# Durable alveolar engraftment of PSC-derived lung epithelial cells into immunocompetent mice

**DOI:** 10.1101/2022.07.26.501591

**Authors:** Michael J Herriges, Maria Yampolskaya, Bibek R Thapa, Jonathan Lindstrom-Vautrin, Feiya Wang, Cheng-Lun Na, Liang Ma, McKenna M Montminy, Jessie Huang, Carlos Villacorta-Martin, Pankaj Mehta, Darrell N Kotton

## Abstract

Durable reconstitution of the injured distal lung epithelium with pluripotent stem cell (PSC) derivatives, if realized, would represent a promising potential therapy for diseases that result from alveolar damage. Here we differentiate murine PSCs in vitro into self-renewing lung epithelial progenitors able to engraft into the injured distal lung epithelium of immunocompetent, syngeneic mouse recipients. Emulating the roadmap of the developing embryo, we generate transplantable PSC-derived Nkx2-1+/Sox9+ lung epithelial progenitors that are highly similar to cultured primary embryonic distal lung bud tip progenitors. These cells display a stable phenotype after frozen archiving or extensive expansion in culture, providing a nearly inexhaustible source of cells that can be engrafted into syngeneic injured mouse lungs without the need for immunosuppression. After transplantation PSC-derived tip-like progenitors downregulate Sox9 and mature in the distal lung, upregulating alveolar type 2 cell markers or assuming the flat morphology and molecular phenotype of terminally differentiated alveolar type 1 cells. After months in vivo, donor-derived cells retain their alveolar epithelial type 2-like and type 1-like phenotypes, as characterized by single cell RNA sequencing, ultrastructural analyses, in vivo histologic profiling, and ex vivo organoid assays that demonstrate continued capacity of the engrafted cells to proliferate and differentiate. These results indicate durable reconstitution of the distal lung’s facultative progenitor and differentiated epithelial cell compartments in vivo with PSC-derived cells, thus establishing a novel model for pulmonary cell therapy which can be utilized to better understand the mechanisms and utility of engraftment prior to future clinical studies.

## Introduction

Acute injuries to the distal lung epithelium, such as those resulting from COVID-19, and chronic lung diseases, such as pulmonary fibrosis or emphysema, represent leading causes of morbidity and mortality worldwide. Common to each of these illnesses is a progressive destruction of the distal lung epithelium that can lead to a lethal reduction in respiratory function. While lung transplants can be used to alleviate symptoms, this solution is severely limited by the insufficient supply of donor lungs and the continual risk of immune rejection of donor tissue despite life-long immunosuppressing drug regimens. One conceivable alternative to full organ transplantation is reconstitution of the injured epithelium through cell therapy, in which donor cells are engrafted directly into a patient to functionally replace lost endogenous cells. While cell therapy has successfully been used to replace bone marrow-derived hematopoietic cells, skin, dopaminergic neurons, dental pulp, corneas, and retinas in patients (Kinoshita et al., 2018; O’Connor et al., 1981; Rama et al., 2010; Schwartz et al., 2015; Schweitzer et al., 2020; Thomas et al., 1957; Xuan et al., 2018), lung epithelial reconstitution in humans has not yet been accomplished. Recent work has shown that cell transplantation is possible in injured mouse lungs with donor-derived cells surviving in vivo and expressing markers of mature epithelial lineages (Kathiriya et al., 2020, 2022; Liao et al., 2022; Louie et al., 2022; Miller et al., 2018; Nichane et al., 2017; Rosen et al., 2015; Vaughan et al., 2015; Xi et al., 2017). However, most of these studies followed the surviving cells for only brief periods in vivo (days to weeks) and utilized either primary lung epithelial cells or immunocompromised mouse recipients, limiting their future potential for clinical application as a cell-based treatment of human lung injury or disease. Furthermore, in many of these studies it is still unclear, with one notable exception (Louie et al., 2022), how donor-derived cells compare to endogenous cells on a wider transcriptional or functional level, which is a critical step towards developing truly therapeutic cell engraftment.

Within the hematopoietic system, similar clinical hurdles and biological questions of progenitor function and perdurance were iteratively solved through mouse models of blood repopulation based on transplantation of mouse hematopoietic progenitor cells into immunocompetent syngeneic recipients, leading to human bone marrow transplant and peripheral blood stem cell transplant therapies that are now standard-of-care for a variety of blood diseases worldwide (Boieri et al., 2016; Morgan et al., 2017). Development of a similar syngeneic murine transplantation assay for the lung epithelium has the potential to provide insight into the treatment and regeneration of this organ and can inform future work in human pre-clinical studies. However, a clinically relevant source of engraftable progenitors for the distal alveolar lung epithelium is not readily apparent since alveolar type 2 (AT2) cells, the endogenous progenitors of this tissue, are difficult to access from most patients and are not easily expanded in cell cultures for autologous therapy (Alysandratos et al., 2021, 2022). Cultured primary cells isolated from the murine distal fetal lung bud tip (hereafter referred to as primary tip-like cells) have been transplanted previously into alveoli (Nichane et al., 2017), making these developmental progenitors a compelling source of donor cells, but a similar human population of autologous embryonic tip cells would be difficult to acquire. Pluripotent stem cell (PSC)-derived cells represent a promising population for syngeneic transplantation, since they provide solutions to these hurdles. For example, using well established protocols, mouse or human induced PSCs (iPSCs) can be generated from any individual without the invasive procedures needed to collect primary distal lung progenitors (Somers et al., 2010; Takahashi and Yamanaka, 2006; Takahashi et al., 2007). In the case of patients with genetic disorders, CRISPR gene editing can then be used to reverse disease-causing mutations, creating a gene-corrected, syngeneic, freezable, and expandable population of cells for generating differentiated donor cells of pulmonary lineages (Alysandratos et al., 2021).

Here we present the derivation and in vivo engraftment of mouse PSC-derived alveolar epithelial progenitors that can durably reconstitute the injured distal lung epithelium of immunocompetent, syngeneic recipient mice. We first generate donor cells by developing a protocol for the directed differentiation of PSCs in order to produce distal lung epithelial progenitors that are transcriptionally similar to transplantable cultured primary tip-like cells (Nichane et al., 2017). When transplanted into bleomycin-injured lungs, these PSC-derived cells integrate into the endogenous alveolar epithelium, reconstituting the desired facultative progenitor function to produce alveolar epithelial type 2 (AT2)-like and alveolar epithelial type 1 (AT1)-like cells. Importantly, these donor-derived cells can persist for at least 6 months in an immunocompetent host and feature functional AT2-specific organelles, such as lamellar bodies. These results demonstrate successful engraftment of PSC-derived cells into an immunocompetent host and provide an important guidepost for developing clinically relevant PSC-derived pulmonary cell therapy without the need for immunosuppression.

## Results

### Lung epithelial specification

In order to generate PSC-derived tip-like cells for transplantation we sought to emulate pulmonary development in which anterior foregut endoderm is specified into early Nkx2-1+ primordial lung progenitors (Ikonomou et al., 2019), which later give rise to fetal distal lung bud tip cells, the developmental precursors of bronchial and alveolar epithelia (Rawlins et al., 2009). Since Nkx2-1 is expressed by all known lung epithelia (Ikonomou et al., 2019), we used a mouse embryonic stem cell (ESC) line with an mCherry reporter targeted to the 3’UTR of the endogenous Nkx2-1 locus (hereafter Nkx2-1^mCherry^) to track, quantify, and purify putative ESC-derived lung epithelial cells (Bilodeau et al., 2014; Serra et al., 2017). To provide an initial basis for our lung specification protocol, we used our previously published directed differentiation approach (Ikonomou et al., 2019) stimulating ESC-derived foregut cells with WNT3a and BMP4 to induce Nkx2-1^mCherry^ expression in a subset of cells (WB protocol, Fig. 1A). Recent single-cell RNA sequencing (scRNA-seq) of murine lung specification in vivo has verified WNT and BMP pathways are active throughout the period of lung lineage specification, but also revealed a switch from retinoic acid (RA) to Fgf signaling soon after Nkx2-1 is first expressed, around E9.5 (Han et al., 2020). To recapitulate this switch in vitro, we added supplemental RA to our specification media until day 8 (D8), when Nkx2-1^mCherry^ is still barely detectable (Fig. 1B), and then added rmFGF10 after D8 (WBRF protocol, Fig. 1A). This novel WBRF protocol resulted in a significant increase in the frequency of NKX2-1^mCherry^+ epithelial cells by D14 (53.95 +/- 3.84%) compared to either the original WB protocol (Ikonomou et al., 2019) or the addition of either supplement alone (Fig. 1B-D). WBRF increased both the percent of epithelial cells that were NKX2-1^mCherry^+ and the overall yield of NKX2-1^mCherry^+/EpCAM+ double positive putative lung epithelial progenitors (Fig. 1E).

**Figure 1:**
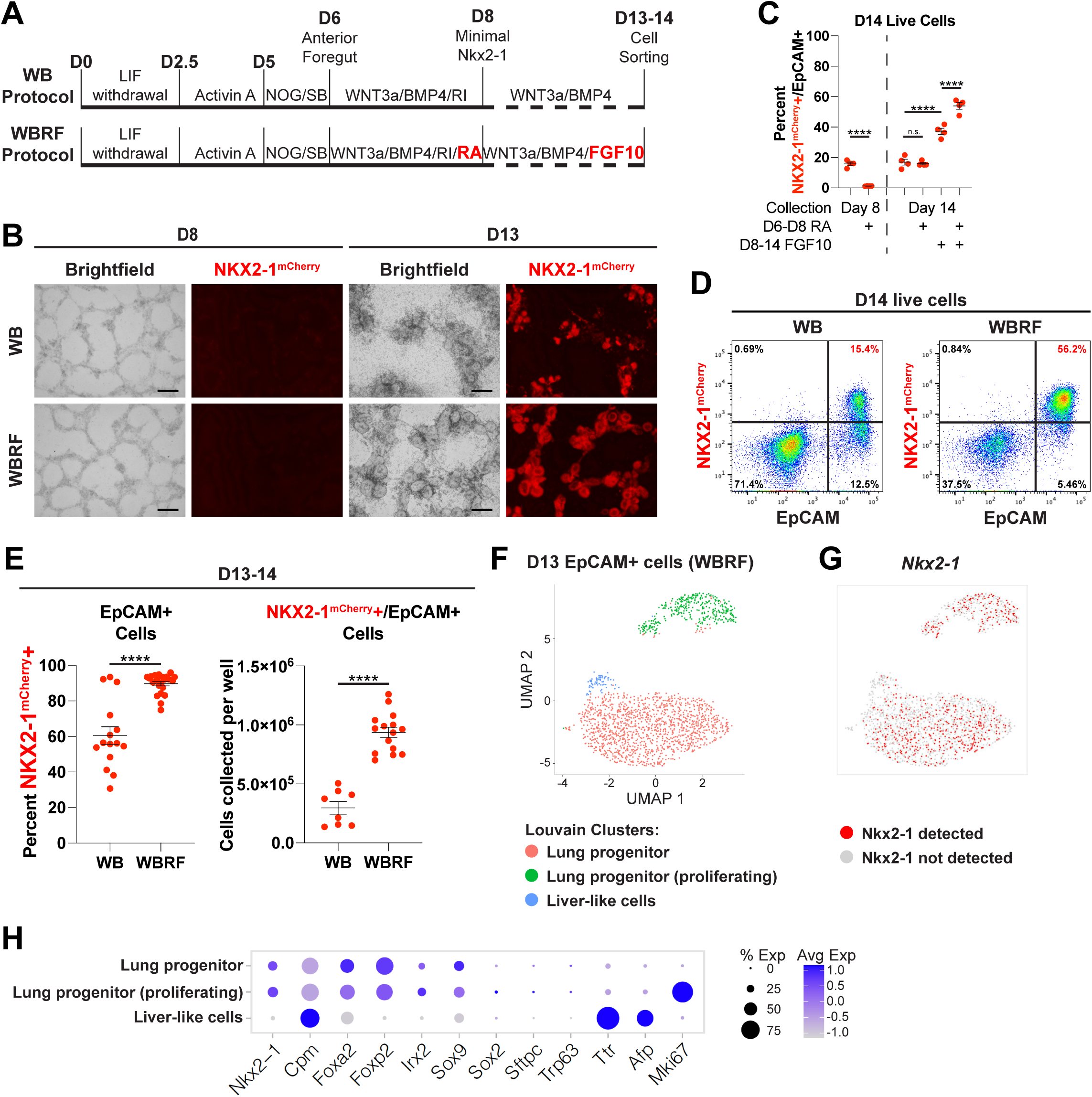
Modified Lung Specification Protocol Leads to Increased Yield of Lung Progenitors. (A) Schematic of WB and WBRF lung specification protocols highlighting the addition of RA and Fgf10 in the WBRF protocol for directed differentiation of mouse ESCs carrying an Nkx2-1^mCherry^ reporter. (B) Representative images of cells during day 8 and day 13 of WB and WBRF lung specification protocols. Scale bars are 200um. Red=mCherry fluorescence microscopy. (C) Quantification of the percent of live cells that are NKX2-1^mCherry^+/EpCAM+ double positive lung epithelial progenitors on day 8 or day 14 depending on the inclusion of RA and/or FGF10. n.s. not significant, **** p < 0.0001 by one-way ANOVA. n=4 biological replicates. Dots with error bars = average +/- SEM. (D) Representative FACS plots on day 14 of WB and WBRF lung specification protocols. (E) Quantification of the percent of epithelial cells that are NKX2-1^mCherry^+ and the yield of NKX2-1^mCherry^+/EpCAM+ double positive cells on day 13-14 of lung specification protocols. **** p < 0.0001 by unpaired, two-tailed Student’s t-test. n=15, 22, 8, 15 biological replicates. Dots with error bars = average +/- SEM. (F) UMAP plot representing scRNA-seq of Epcam+ sorted day 13 cells from WBRF lung specification protocol. Plot displays three clusters identified by Louvain clustering. (G) UMAP plot displaying presence or absence of detectable *Nkx2-1* in these same day 13 cells. (H) Expression of select pulmonary and liver genes in annotated clusters of scRNA-seq data set. See also figure S1.

Consistent with their putative primordial progenitor state (Ikonomou et al., 2019) soon after lung lineage specification and prior to further differentiation, sorted Nkx2-1^mCherry^+/EpCAM+ cells from the WB and WBRF protocols expressed similar levels of early lung epithelial marker transcripts (*Nkx2-1, Foxp2*) and progenitor markers (*Sox9, Id2, Sox2*) (RT-qPCR; Fig. S1A). Distal differentiation markers (*Sftpc* and *Etv5*) were expressed at similar but low levels in cells from each protocol, further emphasizing their primordial state. Both protocols produced cells expressing only low levels of non-lung epithelial lineage markers, with WB inducing low but detectable expression of the thyroid marker *Pax8* (CT>30) and WBRF inducing expression of liver markers (*Afp, Alb*). Finally, low level expression of basal cell markers (*Trp63, Krt5*) was present in cells grown in WB, but not those grown in WBRF. This may suggest that WBRF results in reduced proximal airway fate capacity, consistent with the known distalizing role of FGF10 during lung development (Cardoso and Lü, 2006). To test the lung differentiation competence of each specified population we plated D13 NKX2-1^mCherry^+/EpCAM+ double positive cells from WB and WBRF protocols in our previously published culture conditions that promote expression of distal alveolar or proximal airway epithelial lineage markers (Fig. S1B-D) (Longmire et al., 2012; McCauley et al., 2017). Cells from either specification protocol maintained a high percentage of NKX2-1^mCherry^+ cells in distalizing conditions and expressed similar levels of distal markers (*Sox9, Ager, Sftpc*). Alternatively, proximalizing conditions induced expression of proximal markers (*Sox2, Trp63, Krt5, Scgb3a2*) in both cell populations (Fig. S1E), but as expected WBRF-specified cells gave rise to fewer NKX2-1^mCherry^+ cells with lower expression of these markers. Thus, while cells specified in either protocol are competent to upregulate both airway and alveolar markers, cells specified in WBRF have a reduced efficiency for proximal airway differentiation.

To better understand the heterogeneity of cells generated through the WBRF protocol we profiled all live EpCAM+ cells on D13 by scRNA-seq. Uniform manifold approximation and projection (UMAP) analysis revealed the vast majority of cells localized to two *Nkx2-1*+ clusters, which were predominantly distinguished by expression of cell cycle genes (Fig. 1F, G, S1F). Cells in these two clusters also expressed other primordial lung epithelial associated transcripts (*Cpm, Foxa2, Foxp2, Irx2, Sox9*), but featured minimal expression of proximal marker, *Sox2,* or more differentiated lineage markers (*Sftpc, Trp63*) (Fig. 1H). In line with these results, immunohistochemistry revealed that D13 epithelial cells express nearly ubiquitous NKX2-1, low but detectable levels of the distal progenitor marker SOX9, and no detectable proSFTPC (Fig. S1G). In addition to the two lung lineage clusters, a minor third cell cluster (4.07% of cells) expressed liver markers (*Ttr, Afp*) and minimal *Nkx2-1*. This cluster likely represents the small percentage of NKX2-1^mCherry^-/EpCAM+ non-lung endodermal cells expected in culture prior to any lineage-specific cell sorting (Fig. 1D,1E) (Hurley et al., 2020; Ikonomou et al., 2019). Altogether this data indicates that the novel WBRF protocol can efficiently generate a population of early NKX2-1+ lung epithelial progenitors competent to subsequently differentiate toward airway or alveolar fates.

### Generation of ESC-derived tip-like progenitor cells

Having generated an early lung epithelial progenitor population, we next sought to differentiate these cells into tip-like cells for pulmonary cell transplantation. To do this, we plated D14 NKX2-1^mCherry^+/EpCAM+ double positive cells in Lung Progenitor Media (LPM) culture conditions, similar to those published for maintenance of the progenitor state of murine primary embryonic day 12.5 (E12.5) tip cells (Fig. 2A) (Nichane et al., 2017). In parallel, we generated primary control lines through culturing E12.5 lung epithelial cells from syngeneic mice (hereafter referred to as 129X1/S1) in identical LPM conditions. Both ESC-derived and primary cells grew out as hollow monolayered epithelial spheres that could be passaged multiple times without losing their proliferative capacity or accumulating karyotypic abnormalities (Fig. 2B,C, and S2). ESC-derived tip-like cells could be frozen down and thawed for later use, similar to primary tipl-like cells (Nichane et al., 2017). Furthermore, passaged cells maintained their lung lineage identity, as indicated by retained expression of both the Nkx2-1^mCherry^ reporter and NKX2-1 nuclear protein (Fig. 2D,E). In both the primary and ESC-derived cells we saw a mixture of SOX9^High^/proSFTPC^Low^ and SOX9^Low^/proSFTPC^High^ spheres, suggesting some heterogeneity in alveolar epithelial maturation in LPM conditions (Fig. 2E).

**Figure 2:**
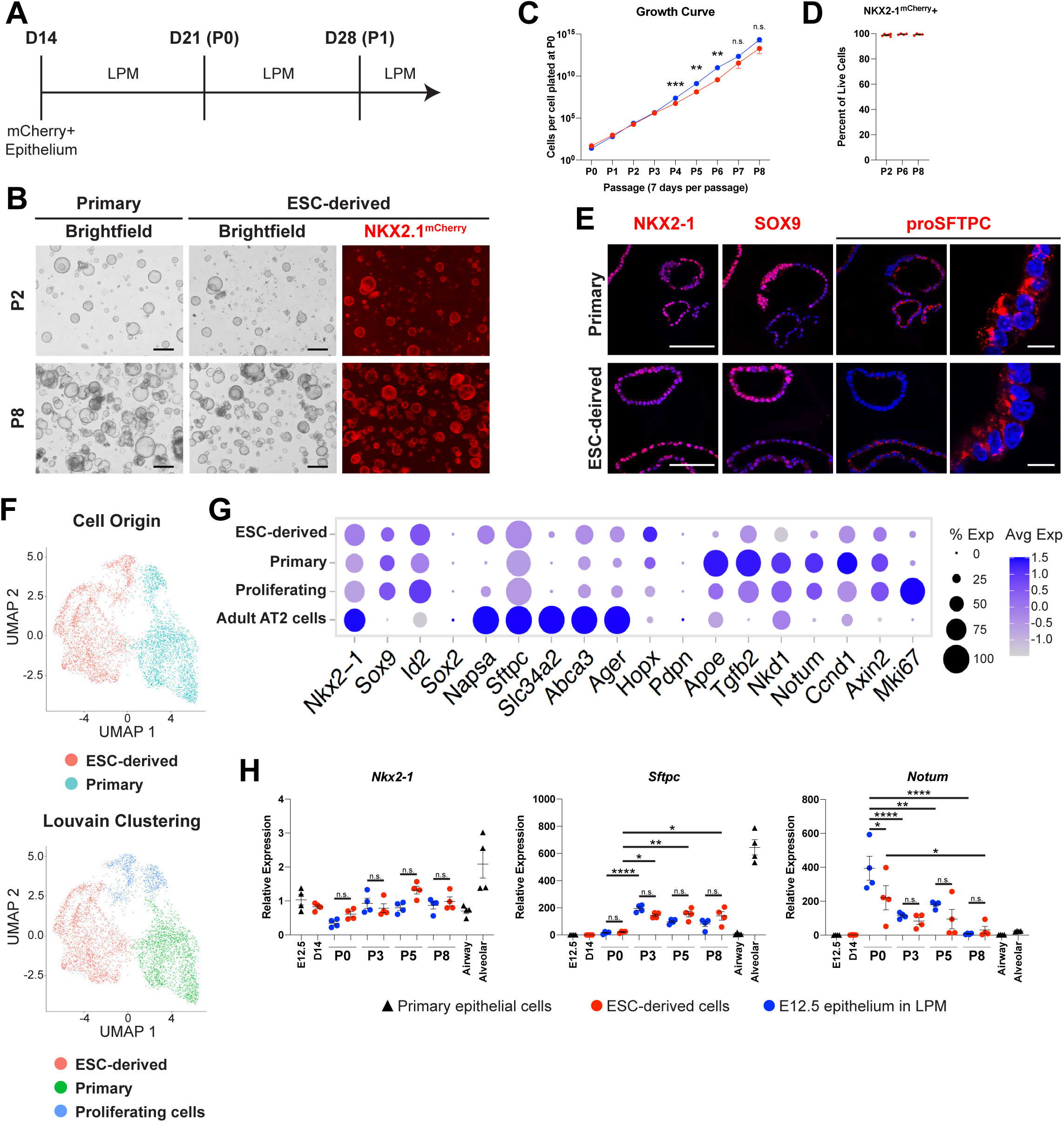
ESC-derived Tip-like Cells are Morphologically and Transcriptionally Similar to Cultured Primary Tip-like Cells. (A) Schematic of differentiation and passaging of ESC-derived tip-like cells in lung progenitor medium (LPM) following cell sorting of Nkx2-1^mCherry+^ cells on day 14. One passage (P) lasts for seven days. (B) Representative images of primary and ESC-derived tip-like cells at P2 and P8. Scale bars are 500um. Red=fluorescence microscopy of the mCherry reporter targeted to the Nkx2- 1 locus. (C) Quantification of cell proliferation for primary and ESC-derived tip-like cells across nine passages. n.s. not significant, ** p<0.01, *** p<0.001 by unpaired, two-tailed Student’s t- test. n= 4 biological replicates. Dot with error bars= average +/- SEM (D) Assessment of NKX2-1^mCherry^ expression throughout passaging of ESC-derived tip-like cells. n= 4 biological replicates. Dots with error bars = average +/- SEM. (E) Immunofluorescence microscopy for NKX2-1, SOX9, and proSFTPC in primary and ESC- derived tip-like cells. Scale bars are 100um (leftmost panels) or 10um (rightmost panels). (F) UMAP plot representing scRNA-seq of primary and ESC-derived tip-like cells. Top plot distinguishes cells by sample origin; bottom plot displays Louvain clusters. (G) Expression of genes, including those identified as differently expressed between primary and ESC-derived tip-like cells. Cells from clusters annotated in panel F are compared against primary adult AT2 cells collected and sequenced at the same time. (I) Analysis of gene expression by RT-qPCR. Primary and ESC-derived tip-like cells from multiple passages are compared against lung epithelial progenitors from day 14 of the WBRF protocol (D14) and freshly sorted lung epithelial cells from embryonic (E12.5) and adult (Airway and Alveolar) mouse lungs. Reference gating for primary controls can be found in supplemental figure 3C,D. n.s. not significant, * p<0.05, ** p<0.01, **** p<0.0001 by one-way ANOVA. n= 4 biological replicates. Dots with error bars = average +/- SEM. See also figures S2-S3.

We then performed single-cell RNA-sequencing (scRNA-seq) of primary and ESC-derived cells from parallel cultures in LPM conditions seven days post-passaging to compare gene expression within these two populations. UMAP visualization of their global transcriptomes (Fig. 2F) with Louvain clustering analysis indicated that the cells segregated partially based on their sample of origin, forming two major clusters, plus a third cluster of actively proliferating cells from both samples. These datasets confirmed similar expression of tip cell markers (*Sox9, Id2*) in cells of all clusters, with relatively low expression of mature AT2 markers, indicating the vast majority of cells of either ESC or primary cell origin were tip-like cells (Fig. 2G). Analysis of differentially expressed genes (DEGs) between the primary and ESC-derived tip-like cells indicated that the ESC-derived cells had higher expression of genes associated with AT2 cells and surfactant metabolism (Fig. S3A,B, Table S1, S2). However, these genes were still expressed at levels well below those of mature primary AT2 cell controls collected at the same time (Fig 2G). On the other hand, primary tip-like cells had higher expression of genes involved in WNT and TGFB signaling (Fig. S3A,B).

To verify these results and screen for any potential drift in gene expression over serial passaging in LPM, we performed RT-qPCR on passage 0 (P0), P3, P5, and P8 of primary and ESC-derived tip-like cells and compared these samples against D14 cells, freshly collected primary tip cells, adult airway epithelial cells, and adult alveolar epithelial cells (Fig. 2H, S3C-E). Lung epithelial progenitor markers (*Nkx2-1, Sox9, Sox2*) were expressed at similar levels in both primary and ESC-derived tip-like cells without alteration after passaging. There were no consistent significant gene expression differences between passage-matched primary and ESC-derived cells for any of the AT2 markers analyzed (*Sftpc, Abca3, Sftpb, Lamp3, Slc34a2, Napsa*). While *Sftpc* and *Abca3* expression did increase from P0 to P3, these increases did not differ based on cell of origin and there were no further increases after subsequent passaging through P8. Similarly, we observed no consistent differential expression based on cell of origin for other selected genes by RT-qPCR (*Notum, Nkd1, Axin2, Tgfb2, Apoe*), but these genes seemed to steadily decline over the first few passages (Fig. S3E). Altogether, this suggests that ESC-derived tip-like cells are transcriptionally similar to primary tip-like cells and maintain their progenitor profile even after expansion in cell culture over multiple passages.

### Transplantation of primary tip-like cells into immunocompetent recipients

One of the ultimate goals of cell therapy is the transplantation of syngeneic cells, alleviating the need for immunosuppression. While primary tip-like cells have been successfully transplanted into NOD-SCID *Il2rg^-/-^*(NSG) mice, it is still unclear whether these transplants can survive in an immunocompetent recipient (Nichane et al., 2017). To test this, we generated primary tip-like cells from UBI-GFP C57BL/6 mice with ubiquitous GFP expression, thus enabling tracking of donor-derived cells following transplantation. Syngeneic C57BL/6J recipient mice were given bleomycin intratracheally to injure endogenous alveolar epithelial cells (Fig. S4A). Ten days later, 6e5 cultured primary tip-like cells were intratracheally instilled. GFP+ donor-derived cells were detected in recipient distal lungs at 9 weeks post-transplantation, indicating long term survival of transplanted cells. These cells appeared in alveolar regions as cuboidal cells with punctate proSFTPC protein immunostaining, characteristic of AT2 cells, as well as thin cells expressing PDPN, characteristic of AT1 cells (Fig. S4B white arrowheads and yellow arrows, respectively). These results suggest that primary tip-like cell transplants can survive and differentiate in syngeneic immunocompetent recipients, similar to previously published transplants into NSG recipients (Nichane et al., 2017).

### ESC-derived tip-like cells give rise to persistent AT2- and AT1-like cells following transplantation

Given the transcriptional similarity between ESC-derived and primary tip-like cells, we next sought to determine whether ESC-derived cells could also be transplanted into immunocompetent mouse lungs. ESC-derived tip-like cells were first labeled with lentiviral GFP for donor cell tracking (Fig. 3A). While these cells were sorted to enrich for GFP+ cells, not all cells maintained GFP expression, likely due to lentiviral silencing (Fig. 3B). Syngeneic 129X1/S1 recipient mice were injured with bleomycin and received 5e5-7e5 donor cells, delivered intratracheally ten days later (Fig. 3C). At three days post-transplantation we observed small, scattered clusters of donor-derived cells (Fig. S5). These cuboidal cells included SOX9+/MKI67+ and proSFTPC+/MKI67+ cells, suggesting they were still in a proliferative tip-like progenitor state. By two weeks post-transplantation, these transplants gave rise to larger clusters of NKX2-1+ cells (Fig. S6A) indicating maintenance of lung epithelial identity. However, few of these cells were SOX9+ or MKI67+ (Fig. S6B, white arrowhead), suggesting loss of the tip-like progenitor identity and reduced proliferation. These donor-derived clusters instead contained both cuboidal cells with punctate proSFTPC and thin PDPN+ cells, suggesting differentiation into AT2-like and AT1-like cells, respectively (Fig. S6C, white arrowheads and yellow arrows). Notably, a subset of donor-derived clusters featured thin AT1-like cells that were largely PDPN-, suggesting incomplete AT1 maturation (Fig. S6C, blue triangles). Altogether this suggests that within 2 weeks, transplanted cells quickly progress from proliferating tip-like progenitors to AT2-like and AT1-like cells, but the degree of maturation may be variable between donor-derived clusters at this early timepoint.

**Figure 3:**
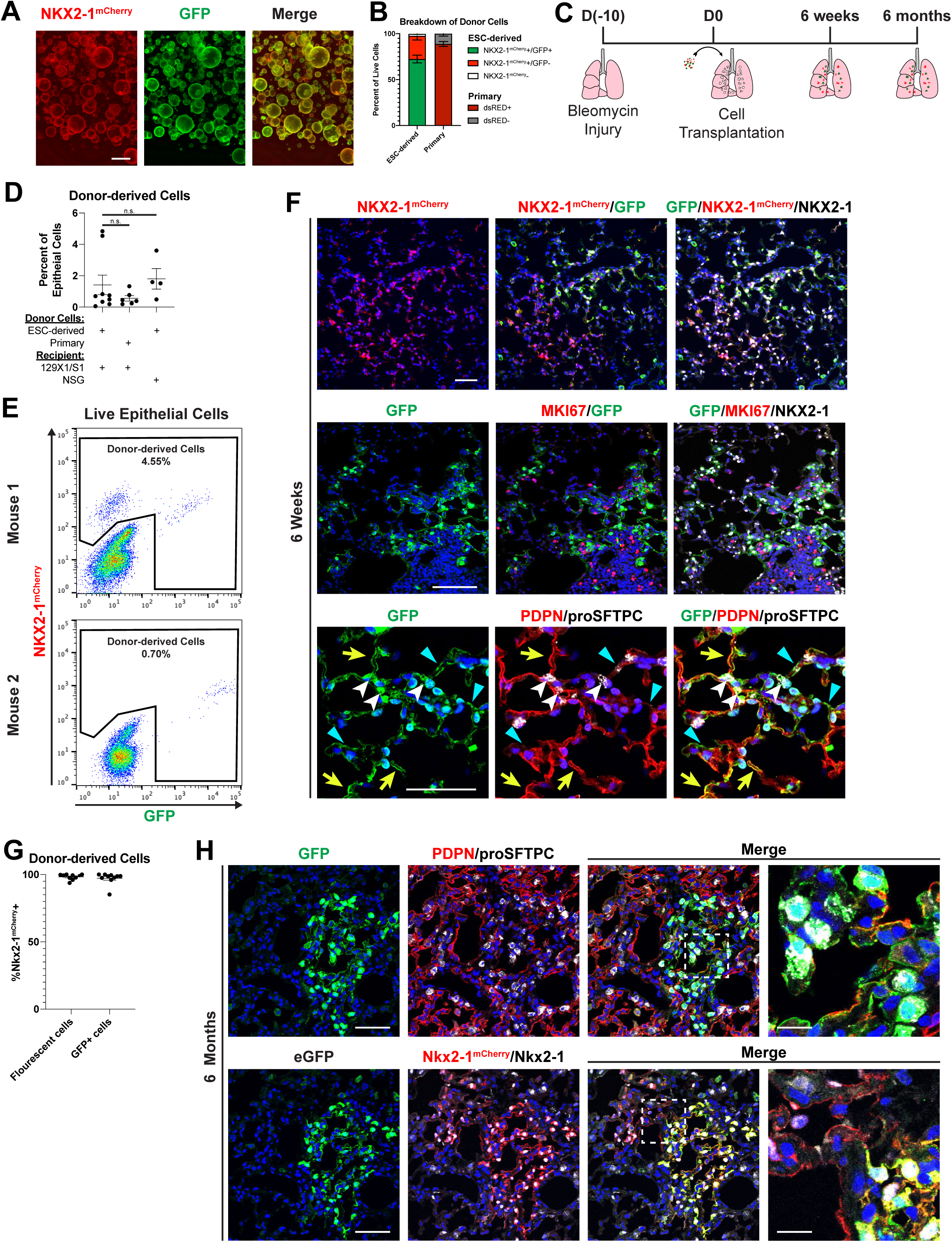
Transplanted ESC-derived Tip-like Cells Give Rise to AT2-like and AT1-like that Persist for At Least Six Months Post-transplantation in Immunocompetent Mice. (A) Representative image of ESC-derived tip-like cells, carrying an Nkx2-1^mCherry^ reporter (red), labeled with lentiviral GFP (green). Scale bar is 500um. (B) Percentage of NKX2-1^mcherry^+/GFP+, NKX2-1^mcherry^+/GFP-, and NKX2-1^mcherry^- cells ESC-derived tip-like cells prior to transplantation (n=5 distinct lines). Also shown is the average dsRed+ percentage of cultured primary cells similarly labeled with lentiviral dsRed (n=3 technical replicates of the same line. Note: primary cells do not have an Nkx2-1 reporter. Error bars = average +/- SEM (C) Schematic for transplantation of cells into bleomycin injured immunocompetent lungs with later histological or flow analysis of donor-derived cells. (D) Flow cytometry quantitation of the percent of live epithelial (EpCAM+) cells that are donor-derived after transplantation of ESC-derived or primary tip-like cells based on flow analysis of recipient lungs. Recipient mice were either immunocompetent 129X1/S1 mice (6 weeks post-transplantation) or immunocompromised NSG mice (9 weeks post-transplantation). n= 9, 6, 4 biological replicates. Dots with error bars = average +/- SEM. (E) Representative FACS plots identifying donor-derived cells within all live lung epithelial cells using mCherry and GFP expression. (F) Representative immunofluorescence confocal microscopy of lung tissue sections showing donor-derived cell clusters at 6 weeks post-transplantation, assessing markers of donor trackers, lung lineages, and proliferation. White arrowheads indicate cuboidal proSFTPC+/GFP+ cells, yellow arrows indicate thin PDPN+/GFP+ cells, and blue triangles indicate thin PDPN-/GFP+ cells. Scale bars are 100um. (G) The percent of all fluorescent (GFP+ or mCherry+) or all GFP+ donor-derived cells that express NKX2-1^mCherry^ at 6 weeks post-transplantation as determined by flow cytometry (for lungs with donor-derived cells accounting for >0.5% of assessed epithelium). n= 10 biological replicates. Dots with error bars = average +/- SEM. (H) Representative histology of donor-derived cell clusters at 6 months post-transplantation assessing markers of donor trackers and lung lineages. Scale bars are 50um (leftmost panels) or 12.5um (rightmost panels). Lower panels indicate some mCherry+ cell clusters are GFP-, presumed due to lentiviral silencing before or after transplantation. See also figures S4-S7

To characterize the durability of donor-derived cells in immunocompetent recipients, we followed mice for longer periods post-transplantation of ESC-derived tip-like cells. By 6 weeks post-transplantation donor-derived cells accounted for 1.4% of all live lung epithelial cells, similar to results seen following transplantation of primary tip-like cells labeled with a lentiviral dsRed into syngeneic 129X1/S1 mice or transplantation of ESC-derived cells into immunocompromised NSG mice (Fig. 3B,D,E). The vast majority of these donor-derived cells were Nkx2-1^mCherry^+/NKX2- 1+/MKI67-, suggesting maintenance of a quiescent lung epithelial fate (Fig. 3F). Flow analysis confirmed that only a small fraction of donor-derived cells were GFP+/Nkx2-1^mCherry^- (2.75% average and 1.2% median, Fig. 3G), indicating that differentiation into non-lung lineages was rare. Similar to two weeks post-transplantation, donor-derived cells included AT2-like cells, AT1-like cells, and thin PDPN- cells (Fig. 3F white arrowheads, yellow arrows, and blue triangles, respectively). Finally, to assess the perdurance of transplanted cells in the presence of a functional immune system, we dissected mice at six months post-transplantation. Even at this late timepoint we were able to find large clusters of donor-derived AT2-like and AT1-like cells (Fig. 3H), suggesting long term survival of donor-derived epithelial lineages. Together these data suggest that ESC-derived cells transplanted into immunocompetent mice can differentiate into AT2-like and AT1-like cells and durably maintain these identities over time.

### Profiling donor-derived and endogenous cells at single-cell resolution

To better characterize the fate of transplanted ESC-derived cells, we profiled recipient lungs by scRNA-seq 6 and 15 weeks post-transplantation. We first collected live epithelial cells (DRAQ7-/EpCAM+/CD45-/CD31-) from an uninjured control as well as the donor-derived (mCherry+ or GFP+) and endogenous (mCherry-/GFP-) epithelium from transplant recipients 6 and 15 weeks post-transplantation (Fig. 4A, S7, S8A). In order to determine whether donor-derived cells contributed to non-epithelial cell types, we collected all mCherry+ or GFP+ cells from our 15 weeks post-transplantation mouse. The resulting datasets were visualized with UMAP and cells were clustered using the Louvain algorithm (Fig. S8B,C). Non-epithelial clusters were then identified based on expression of *Col1a2*, *Pecam1*, or *Ptprc* for mesenchymal, endothelial, or hematopoietic lineages, respectively (Fig. S8D,E). Importantly, out of the 2,092 non-epithelial cells characterized, only one cell expressed mCherry or GFP (Fig. S8A). This cell expressed low levels of both AT2 and macrophage markers, indicating it was likely a donor-derived AT2-like cell being phagocytosed by a macrophage. Altogether this suggests that donor-derived cells primarily give rise to epithelial lineages.

**Figure 4:**
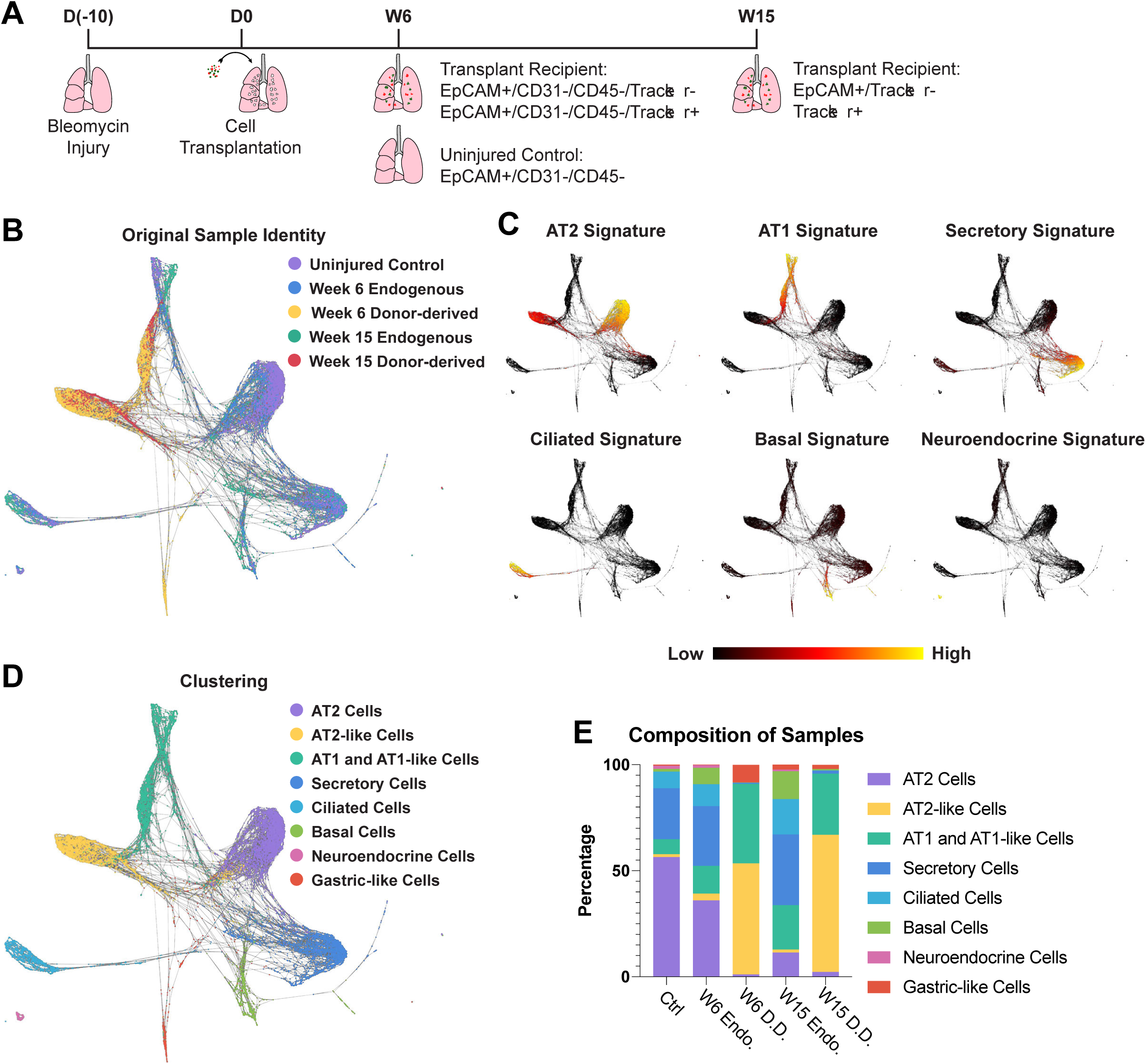
Single Cell Transcriptomic Profiling of Donor-derived and Endogenous Lung Epithelium. (A) Schematic for generation and collection of samples for scRNA-seq. (B) SPRING plot of epithelial cells characterized by scRNA-seq, with cells labeled by sample origin. (C) Expression of lung epithelial cell signatures. Gene sets comprising each signature can be found in Supplementary Table 3. (D) Cell-type annotation of clusters based on supervised Louvain clustering and expression of lung epithelial cell signatures. (E) Composition of each sample based on clusters identified in figure 4D.

**Figure 7:**
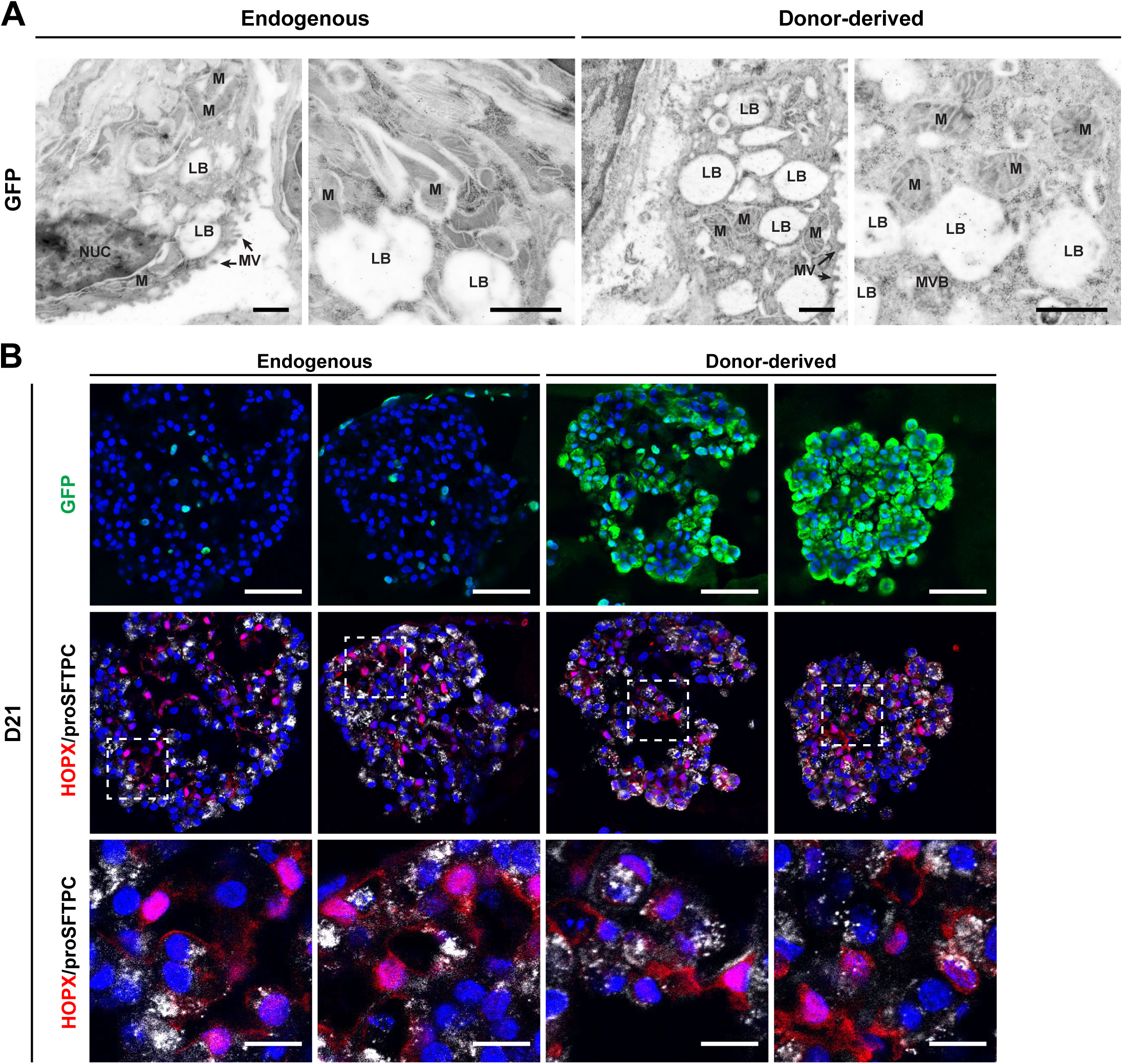
Functional Assessment of Donor-derived AT2-like Cells. (A) Representative transmission electron micrographs of GFP- control AT2 cells and GFP+ donor-derived AT2-like cells with immunogold staining to detect GFP (black dots). Lamellar bodies appear as electron-lucent cytoplasmic areas, rather than lamellated inclusions, due to their extrusion during immunogold processing. Scale bars are 1.0 um. LB lamellar body, NUC nucleus, **M** mitochondria, MV microvilli, MVB multivesicular bodies. (B) Representative immunofluorescence confocal microscopy of GFP/HOPX/proSFTPC expression in cultured mouse lung alveolospheres, comparing endogenous or donor­ derived epithelial cells co-cultured with PDGFRanGFP+ primary lung fibroblasts. PDGFRa­ GFP is nuclear, while donor-derived cells have a cytoplasmic GFP. Scale bars are 50um (top row) or 12.5 um (bottom row). Epithelial cells were sorted from recipient mice 40 days after transplantation.

In order to coalesce the data sets generated from our two transplant recipients, the epithelial cells from each dataset were combined using harmonization prior to plotting with UMAP (Fig. S8F) or combined without harmonization and plotted on SPRING (Fig. 4B) (Korsunsky et al., 2019; Weinreb et al., 2018). The harmonized UMAP dataset was used to divide the cells based on Louvain clustering with overlapping or highly similar clusters being combined manually. The resulting cell clusters were annotated based on expression of lung epithelial cell type gene signatures and included all major endogenous lung epithelial cell types (Fig. 4C,D, S8G, Table S3). Donor-derived cells were predominantly found in three clusters (Fig. 4E). Consistent with immunostaining results (Fig. 3F), the donor-derived samples were mCherry+/GFP+ and did not express any proliferation markers, suggesting they had assumed a quiescent state similar to the endogenous epithelium at this stage (Fig. S9). The vast majority of donor-derived cells expressed high levels of either AT2 or AT1 gene signatures without expressing other cell type signatures (hereafter AT2-like and AT1-like cells, respectively) (Fig. 4C). Finally, a small third cluster from the 6-week post-transplantation sample lacked *Nkx2-1* expression (Fig. S9). These cells (hereafter gastric-like cells) corresponded to the rare mCherry- cells seen in this sample and expressed gastric markers associated with loss of NKX2-1 in lung epithelium (Fig S7A, S9) (Herriges et al., 2017; Snyder et al., 2013). Altogether, this indicates that donor-derived cells primarily give rise to cells transcriptionally similar to endogenous AT2 and AT1 cells.

Although the majority of donor-derived cells expressed alveolar epithelial lineage markers, these cells did not overlap perfectly with endogenous cells, indicating differences in molecular phenotype. In order to identify DEGs for both AT2-like and AT1-like cells, we compared donor-derived and endogenous cells within each of these cell types (Fig. S10A, Table S4, S5). While there was little overlap in DEGs between the two cell types, both donor-derived AT2-like and AT1- like cells were deficient in expression of major histocompatibility complex II (MHC-II) genes (Fig. 5A, Table S6, S7). This deficiency corresponds to a nearly complete absence of MHC-II expression in sterile donor cells prior to transplantation (Fig. S10B). In the adult lung epithelium MHC-II is primarily expressed in AT2 cells, where it contributes to antigen presentation, which in turn regulates resident memory T cell function and barrier immunity during infection (Shenoy et al., 2021; Toulmin et al., 2021). Even though the majority of donor-derived AT2-like cells remained deficient for MHC-II genes even at 15 weeks post-transplantation, a portion of these did express endogenous levels of MHC-II components (Fig. 5B). This suggests that with sufficient time and exposure to a non-sterile environment, donor-derived cells may upregulate MHC-II components.

**Figure 5:**
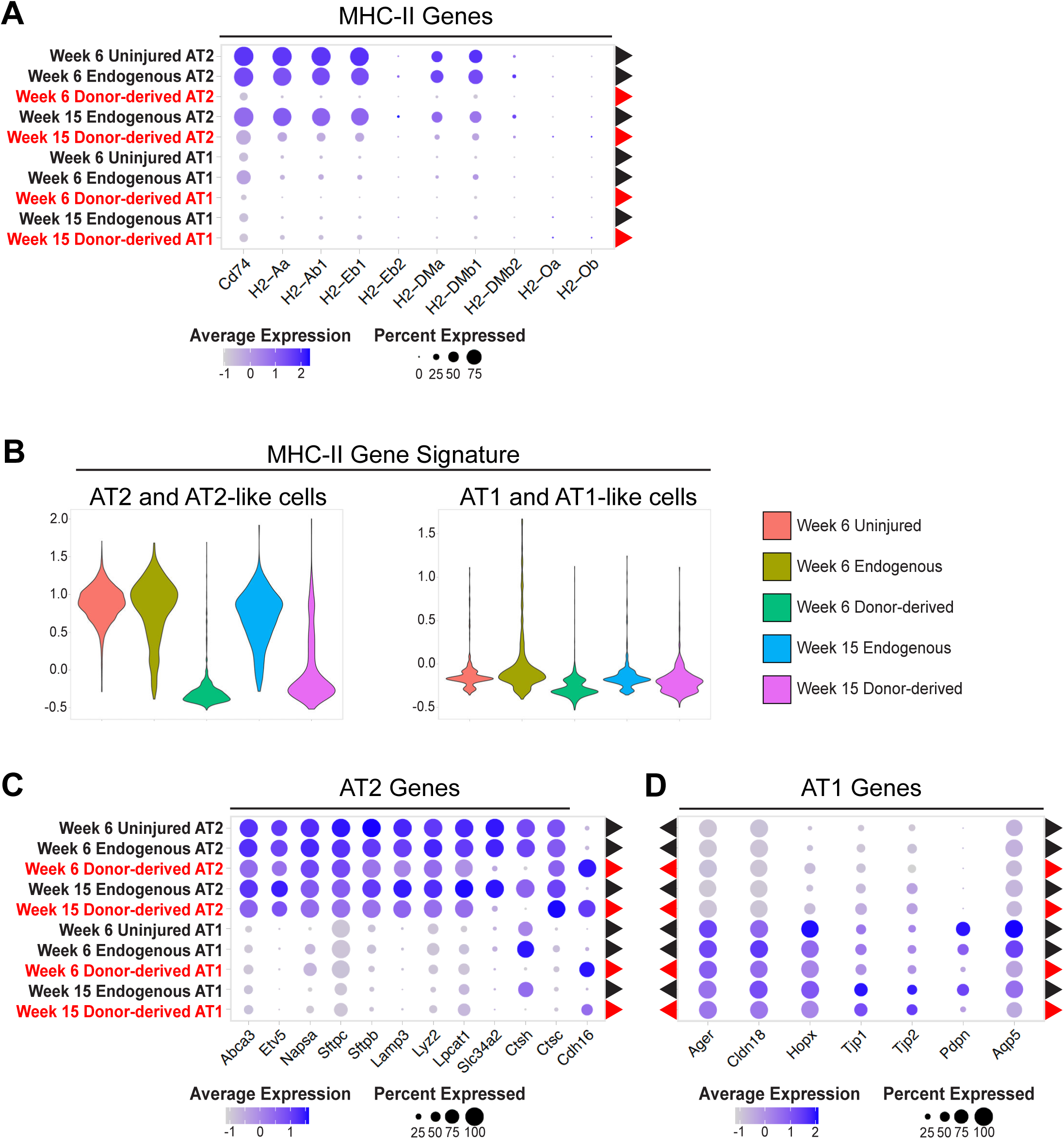
Donor-derived Cells Express Lower Levels of Select MHC-11 Components and Maturation Markers. (A) Expression of MHC-11genes in donor-derived (red) and endogenous (black) cells. (B) Violin plots of an MHC-11gene signature composed of genes listed in figure 5A. (C) Expression of AT2 genes in donor-derived (red) and endogenous (black) cells. (D) Expression of AT1 genes in donor-derived (red) and endogenous (black) cells.

In addition to differences in MHC-II expression, both donor-derived AT2-like and AT1-like cells demonstrated subtle signs of incomplete maturation. Both cell types expressed many of the canonical lineage markers associated with the corresponding endogenous lineage, although some were expressed at lower levels (Fig. 5C,D, S10). Those few markers that were nearly absent in donor-derived cells were associated with late stage maturation of AT2 (*Ctsh, Slc34a2*) or AT1 (*Pdpn, Aqp5*) cells. While CTSH has been shown to play an important role in SFTPB processing, donor-derived cells expressed *Ctsc*, which can compensate for the absence of *Ctsh* (Bühling et al., 2011; Ueno et al., 2004). Few noteworthy genes were significantly upregulated in donor-derived cells, but they did maintain high expression of an embryonic cadherin (*Cdh16*) (Fig. 5C) (Wertz and Herrmann, 1999). Altogether, this indicates that donor-derived cells express many of the genes necessary for alveolar epithelial cell function, but may require further priming before or after transplantation in order to fully mature and respond to the non-sterile environment of the lung.

### scTOP demonstrates global transcriptomic alignment between donor-derived and endogenous alveolar lineages

Differential gene expression analysis, SPRING, and UMAP are all designed to highlight differences in cell populations, but these methods do not provide means to quantify the overall similarity of non-identical cell populations. To provide unbiased quantitative assessments of how well our donor-derived cells align with endogenous alveolar lineages on a global transcriptomic level, we developed a computational algorithm, Single-Cell Type Order Parameters (scTOP) (Fig. 6A and supplemental methods) (Yampolskaya et al., in preparation). While other methods of dimensionality reduction rely on unsupervised machine learning to determine axes of relevance with no prior knowledge, scTOP uses established single-cell atlases as references to determine alignment with known cell types. This algorithm reduces the number of dimensions from the number of genes down to the number of known cell types, retaining more information than methods which reduce the data to 2 dimensions. These reference cell types are used to create vectors, which define the dimensions of a cell type subspace. To find the alignment of a sample of interest with each cell type, and thus locate the sample in cell type space, the transcriptional profile of the sample is projected onto the subspace formed by the reference vectors. This projection can be assessed for either individual cells or the average of a cell population, producing individual or aggregate alignment scores, respectively.

**Figure 6:**
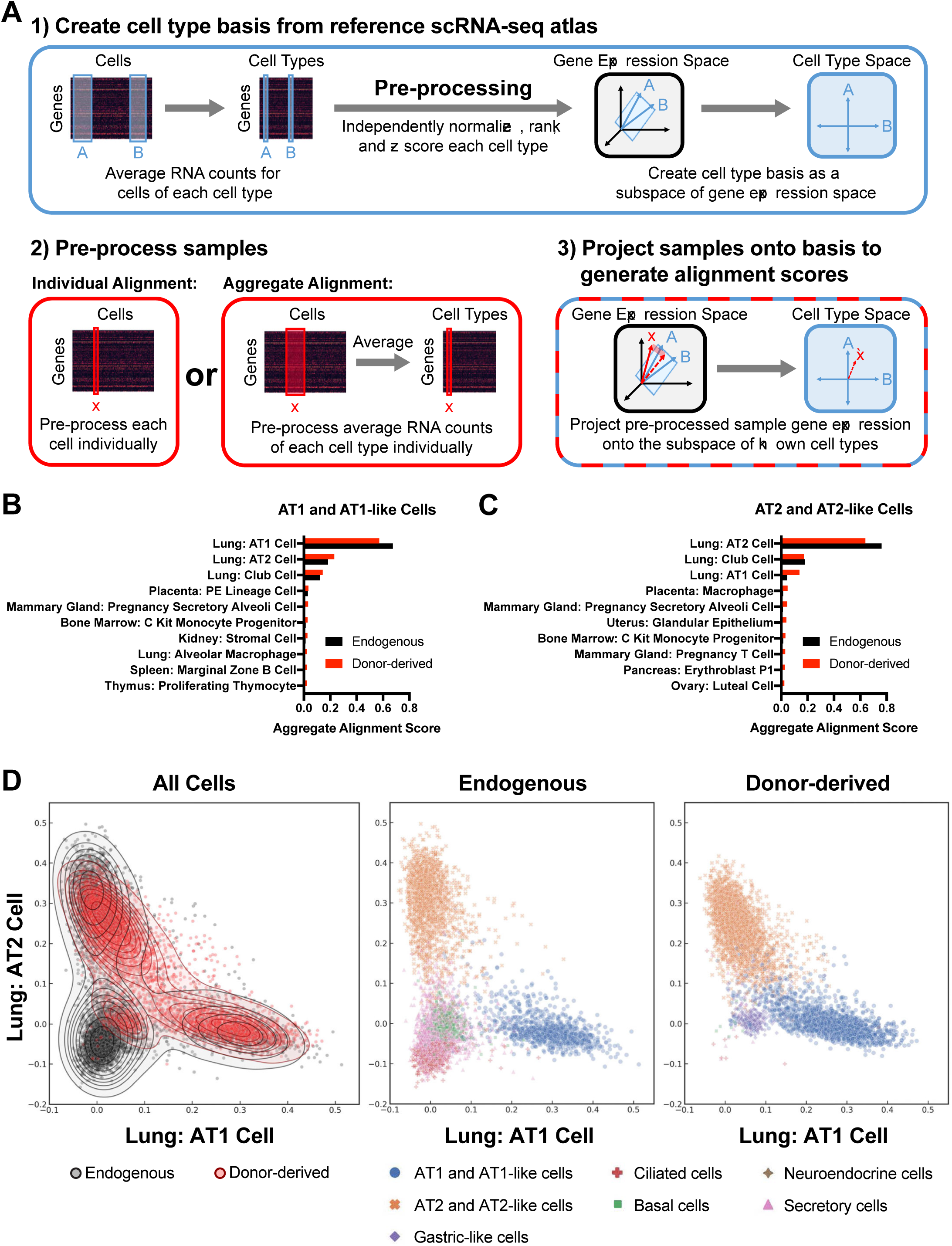
Global Transcriptomic Comparison of Endogenous and Donor-derived Lung Epithelial Cells using scTOP. (A) Schematic of scTOP (Single-Cell Type Order Parameters) used to generate alignment scores. (B) The top ten aggregate alignment scores for donor-derived AT1-like cells and the corresponding scores for endogenous AT1 cells. All reference cell types are from adult mice (Mouse Cell Atlas or Control sample as delineated in Figure 4) (Han et al., 2018). (C) The top ten aggregate alignment scores for donor-derived AT2-like cells and the corresponding scores for endogenous AT2 cells. All reference cell types are from adult mice. (D) Individual alignment scores for all donor-derived and endogenous epithelial cells against reference adult lung AT1 and AT2 cells. Each cell is annotated (by color and shape shown in the key below the graphs) based on sample type or cell type as determined in Figure 4.

In order to determine the alignment scores of our donor-derived and endogenous epithelial cells in transplant recipients, we projected pre-processed cell transcriptomes onto reference data sets compiled from the Mouse Cell Atlas (Han et al., 2018) and the previously described uninjured control mouse lung from figure 4. Donor-derived and endogenous AT1 cells both displayed similar aggregate alignment profiles, with both primarily aligning to the lung AT1 cell reference benchmark with high aggregate alignment scores (0.572 and 0.676, respectively; Fig. 6B). For both populations the next highest alignment was against lung AT2 cells (0.230 and 0.183), potentially reflecting the close lineage relationship between AT1 and AT2 cells. Donor-derived and endogenous AT2 cells also had similar aggregate alignment profiles, with both primarily aligning to lung AT2 cells by a considerable margin (alignment score 0.638 and 0.761, respectively; Fig. 6C). Notably, donor-derived AT2-like cells had higher alignment to lung AT1 like cells compared to endogenously-derived cells (0.138 vs 0.045). Combined with the lower AT2- alignment score, this may reflect either incomplete maturation of donor-derived AT2-like cells or initiation of AT1 differentiation in a subset of this population. In contrast to these alveolar epithelial populations, the rare donor-derived gastric cells did not have a single dominating alignment score, but instead weakly aligned to multiple reference cell types (Fig. S11A). Interestingly, while these cells do not express Nkx2-1, the top four alignment scores are all against lung cell populations, suggesting a maintained lung transcriptional program even in the absence of this critical lung transcription factor.

To assess the individual alignment scores of transplanted cells, we then projected individual cells against the same reference basis (Fig. 6D). In the resulting plots, cells were labeled as either donor-derived or endogenous and color-coded based on their previously assigned lineage identity from figure 4. The vast majority of individual endogenous cells aligned primarily with their cell type predicted in figure 4, confirming our previous identification of these cells (Fig. 6D, S11B,C). As expected, the majority of individual donor-derived cells aligned with either AT1 or AT2 cells, with all gastric-like cells aligning poorly to both cell types. Donor-derived and endogenous AT1 cells had nearly overlapping alignment distributions on all analyzed plots, reflecting substantial transcriptional similarity for these two populations, despite the differentially expressed genes identified above. While endogenous and donor-derived AT2 cells did not overlap as precisely, there was significant overlap between the two distributions suggesting a relatively similar transcriptional profile for at least a subset of cells. As expected based on our lineage marker analysis, few donor-derived cells aligned well with any airway lineage (Fig. S11B,C). Altogether these results indicate that while select genes demarcate donor-derived and endogenous cell populations, the two populations are highly similar and exhibit alveolar cell states when scored on a global transcriptomic level.

### Donor-derived AT2-like are functionally similar to endogenous AT2 cells

In order to establish whether or not transplantation of ESC-derived tip like cells models true cellular engraftment, it is necessary to determine whether donor-derived cells are functionally similar to endogenous alveolar epithelial cells (Alysandratos et al., 2021). The punctate localization of proSFTPC in donor-derived AT2-like cells suggested the presence of lamellar bodies, a secretory organelle critical to AT2 cell secretory function (Fig. 3F). To verify this and characterize the ultrastructural features of GFP+ donor-derived AT2-like cells, we performed transition electron microscopy along with immunogold staining against GFP (Fig. 7A). Donor-derived AT2-like cells were ultrastructurally indistinguishable from AT2 cells in a control mouse and contained lamellar bodies, indicating donor-derived AT2-like cells have the organelles necessary for protein and surfactant secretion.

In addition to their secretory function, AT2 cells are the known facultative progenitors of the adult alveolar epithelium (Adamson and Bowden, 1974; Barkauskas et al., 2013; Evans et al., 1975). In response to injury or cell culture conditions, these normally quiescent cells can proliferate and differentiate into AT1 cells. To determine if donor-derived cells have a similar progenitor function, endogenous or donor-derived epithelial cells were sorted and cultured with PDGFRa^nGFP+^ lung mesenchymal cells (Barkauskas et al., 2013). After 21 days in culture both endogenous and donor-derived cells plated as a single-cell suspension gave rise to large epithelial organoids, indicating that these previously quiescent cells (Fig. 3F) can re-enter the cell cycle (Fig. 7B). Both endogenous and donor-derived cell organoids contained cuboidal proSFTPC+/HOPX- cells as well as proSFTPC-/HOPX+ cells with thin protrusions, suggesting differentiation into AT2-like and AT1-like cells, respectively. In addition to these expected cell types, proSFTPC+/HOPX+ cells could be found in donor-derived organoids. This corresponds to increased *Hopx* expression (Fig. S10) and AT1 alignment (Fig. 6C) in donor-derived AT2-like cells in vivo and potentially indicates instances of delayed or incomplete AT1 differentiation in this co-culture assay. Altogether, this suggests that donor-derived cells can function as facultative progenitors and that the transplants described above model functional cellular engraftment capable of replacing endogenous epithelial cells.

## Discussion

The ultimate goal of syngeneic pulmonary cell therapy is an effective treatment for pulmonary injury and disease that is not dependent on the availability of donor tissue or the use of detrimental immunosuppressants. While previous pioneering studies have demonstrated cell transplantation in mouse lungs, these studies utilized difficult to collect primary cells or necessitated the use of immunocompromised recipients (Kathiriya et al., 2020, 2022; Liao et al., 2022; Louie et al., 2022; Miller et al., 2018; Nichane et al., 2017; Rosen et al., 2015; Vaughan et al., 2015; Xi et al., 2017). To address these issues, we developed an approach to engraft PSC-derived cells into the lungs of syngeneic and immunocompetent recipients. Following transplantation, these cells give rise to persistent AT2-like and AT1-like cells that are transcriptomically and functionally similar to endogenous alveolar epithelial cells. This model thus provides a valuable system to further characterize and optimize pulmonary cell therapy in an approachable and clinically relevant system.

One of the major hurdles of cell transplantation is identifying and producing an effective donor cell population. To create an ESC-derived donor population, we focused on mimicking cultured primary tip-like cells, a cell population which can be transplanted in immunodeficient mice (Nichane et al., 2017). Utilizing this guidepost, along with published differentiation protocols and in vivo development signaling networks (Han et al., 2020; Ikonomou et al., 2019; Serra et al., 2017), we developed a stepwise and efficient protocol for the directed differentiation of mouse ESCs into tip-like epithelial cells. The resulting progenitor population could be frozen down or expanded without losing their cell identity and were transcriptomically similar to cultured primary tip cells without necessitating the collection of primary respiratory tissues. This ability to produce and store large numbers of donor cells will be critical as the field progresses toward clinical cell therapy, where an abundance of cells will likely be needed to treat extensive damage in human lungs.

To assess the viability of syngeneic transplantation, we then transplanted our ESC-derived cells into immunocompetent recipients. Initially these cells maintained their proliferative SOX9+ progenitor identity in vivo. However, as early as 2 weeks post-transplantation the donor-derived cells differentiated into AT2-like and AT1-like cells. Further analysis of this rapid differentiation in future studies will provide important information on the mechanisms behind epithelial repair and guide development of future in vitro differentiation protocols. While multiple papers have assessed pulmonary cell transplant survival within these preliminary weeks post-transplantation, few have assessed the long-term survival of transplanted cells (Louie et al., 2022). Following our ESC-derived transplants we saw differentiated donor-derived cells surviving in vivo for at least 6 months. These cells maintain their alveolar epithelial identity and remain largely quiescent, similar to the endogenous alveolar epithelium. This indicates progenitor cells engineered outside of the lung can differentiate following transplantation and survive for extended periods in the presence of a functional immune system without developing into tumorigenic cells.

PSC-derived cells differentiated in vitro are often substantially different from their mature primary counterparts on a transcriptomic level despite expression of key lineage markers (Abo et al., 2022; Alysandratos et al., 2022; Fidanza et al., 2020; Pavlovic et al., 2018; Schwartz et al., 2014; Xia et al., 2016). Therefore, it was critical to determine whether our engineered cells could differentiate in vivo into cells transcriptomically similar to the endogenous alveolar epithelium. Sc- RNAseq analysis indicated that donor-derived cells were highly similar to the endogenous epithelial lineages with significant expression of many of the canonical markers traditionally associated with these cell types. To provide a more global transcriptomic comparison of endogenous and donor-derived epithelial cells we utilized scTOP, a novel algorithm which allows for quantifiable alignment of individual cells or cell populations against known reference cell types. Through this analysis we observed that our donor-derived lineages had an alignment profile similar to that of paired endogenous cells, which aligned specifically to the expected reference benchmarked cell type. In particular, the alignment of donor-derived AT1-like cells were almost indistinguishable from endogenous AT1 cells. Despite these overall similarities, donor-derived AT2-like and AT1-like cells both had notable deficiencies in expression of select maturation markers and components of the MHC-II complex. Further studies will be needed to understand whether these differences impact cellular function and how they can be addressed to improve transplant performance. In particular, the MHC-II complex represents a highly tractable target for future research. The observed MHC-II deficiency in donor-derived cells may be due to the donor cells originating from a sterile environment with minimal MHC-II expression (Fig. S10B). It is still unclear whether this deficiency simply represents immunological immaturity or whether these differences from endogenous AT2 cells would persist following subsequent lung infection, where antigen presentation by AT2 cells contributes to the immune response (Shenoy et al., 2021; Toulmin et al., 2021). If donor-derived AT2-like cells do not properly respond in vivo, this may indicate a need to stimulate MHC-II expression in donor cells prior to transplantation or use in clinical cell therapy.

Although multiple studies have reported transplantation of cells into the lung, few have gone on to assess the functionality of donor-derived cells (Kathiriya et al., 2020, 2022; Liao et al., 2022; Louie et al., 2022; Miller et al., 2018; Nichane et al., 2017; Rosen et al., 2015; Vaughan et al., 2015; Xi et al., 2017). In order to establish true cellular engraftment and progress towards cell therapy, it is imperative to examine whether donor-derived cells are functionally similar to the target endogenous cells (Alysandratos et al., 2021) In this study we demonstrated that donor-derived AT2-like cells produce lamellar bodies, the specialized organelles necessary for AT2 secretory function. Furthermore, quiescent donor-derived cells were capable of re-entering the cell cycle in culture and producing both AT2-like and AT1-like cells, indicating the existence of facultative progenitors similar to endogenous AT2 cells (Adamson and Bowden, 1974; Barkauskas et al., 2013; Evans et al., 1975).

While this work provides important insights into the feasibility of PSC-based cell therapy, future studies will be needed to further characterize the functionality of engrafted cells and transition towards clinical cell therapy. Our successful transplantation into immunocompetent mice serves as groundwork for modelling cell therapy in existing murine models of pulmonary genetic diseases (Nureki et al., 2018; Rindler et al., 2017). Future work will be needed to functionally test whether engraftment of engineered cells is sufficient to prevent mortality associated with severe progressive pulmonary diseases. In parallel to this work, alternative methods and donor cell populations will be needed to apply lessons learned in the mouse model to the development of clinical pulmonary cell therapies. In particular, while bleomycin is sufficient to clear out some endogenous epithelium, alternative methods of cellular clearance, such as targeted decellularization (Dorrello et al., 2017), may provide more precise and consistent removal of epithelial lineages. Likewise, there will be a need to identify the ideal human iPSC-derived donor lines with current protocols to generate iPSC-derived tip-like or AT2-like cells providing promising candidates (Chen et al., 2017; Huang et al., 2014; Jacob et al., 2017; Miller et al., 2018; Yamamoto et al., 2017).

In summary, the current study establishes a model of PSC-derived pulmonary cell therapy that results in durable engraftment of donor-derived cells in an immunocompetent recipient. This model thus provides an important foundation for further characterization and optimization of pulmonary cell engraftment, with potential to yield important insights into pulmonary regeneration and the development of clinical cell therapies.

---

**Supplemental Figure 1:**
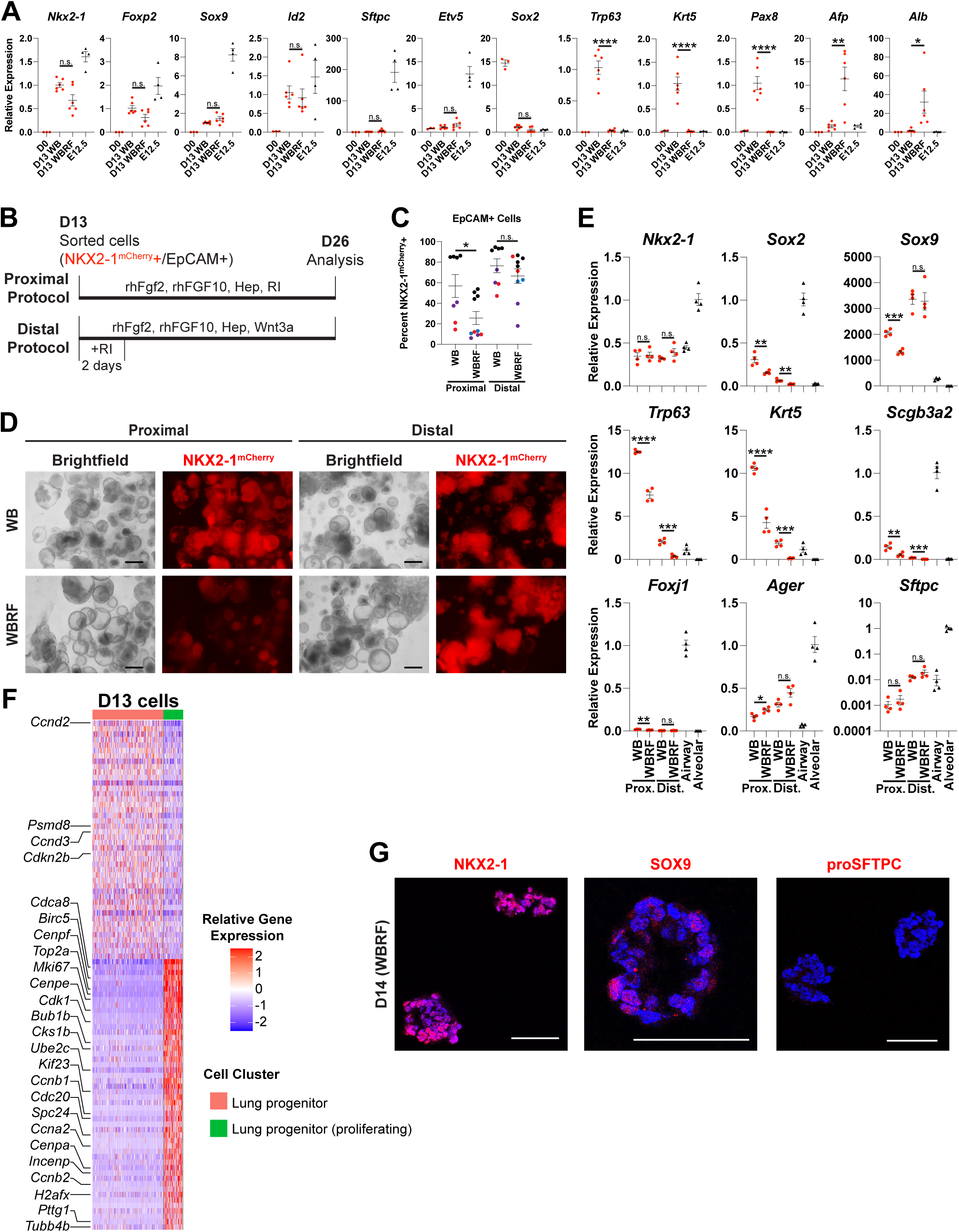
Comparison of Gene Expression and Differentiation Potential for WB-specified and WBRF-specified Lung Epithelial Progenitors. (A) Analysis of gene expression by RT-qPCR at day 13 of WB and WBRF lung specification protocols compared to ESCs (D0) and freshly collected E12.5 SOX9+ epithelium. n.s. not significant, * p<0.05, ** p<0.01, **** p<0.0001 by unpaired, two-tailed Student’s t-test. n= 3, 6, 6, 4 biological replicates. Dots with error bars = average +/- SEM. (B) Schematic for proximal and distal differentiation protocols. (C) The percent of cells that were NKX2-1^mCherry^+ based on specification (WB vs WBRF) and differentiation (Proximal vs Distal) protocols. Dot color indicates biological replicates performed in the same batch. n.s. not significant, * p<0.05 by one-way ANOVA. n= 8, 10, 8, 10 biological replicates. Dots with error bars = average +/- SEM. (D) Representative image of cells on day 26 of proximal and distal differentiation protocols. Scale bars are 500um. (E) RT-qPCR analysis of gene expression in day 26 NKX2-1^mCherry^+ cells of proximal and distal differentiation protocols. Cultured cells were compared to freshly collected epithelial cells from adult mouse lungs (Airway and Alveolar). n.s. not significant, * p<0.05, ** p<0.01, *** p<0.001, **** p<0.0001 by unpaired, two-tailed Student’s t-test. n= 4 biological replicates. Dots with error bars = average +/- SEM. (F) Row-normalized heatmap of the top 50 most up-regulated and top 50 most down- regulated genes (with adj. p-value <0.05, ordered by logFC) between the clusters Lung progenitor and Lung progenitor (proliferating) in day 13 of the WBRF specification protocol. Annotated genes are associated with proliferation. (G) Representative immunofluorescence confocal microsocopy of paraffin tissue sections prepared from day 14 cells cultured in the WBRF specification protocol. Staining performed with antibodies against NKX2-1, SOX9, or proSFTPC. Scale bars are 200um.

**Supplemental Figure 2:**
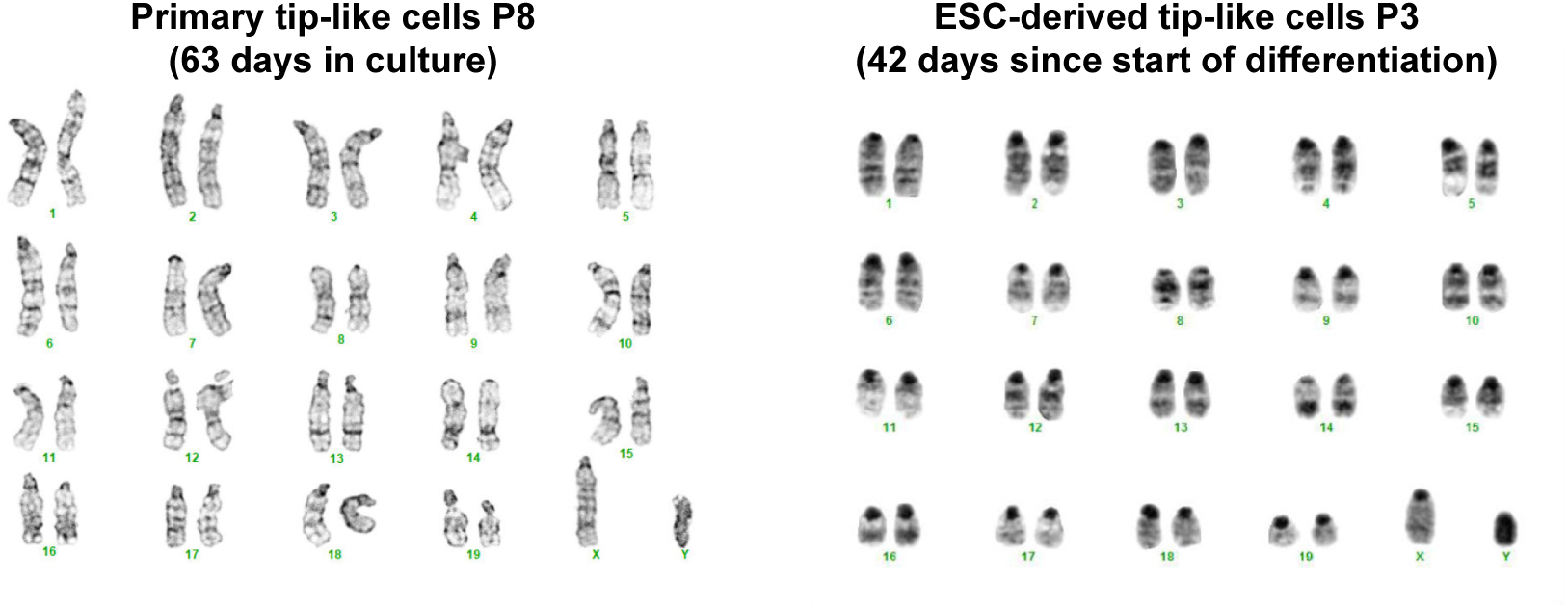
Karyotyping of Passaged Primary and ESC-derived Tip-like Cells. Representative G-banding indicates karyotypically normal primary and ESC-derived tip­ like cells.

**Supplemental Figure 3:**
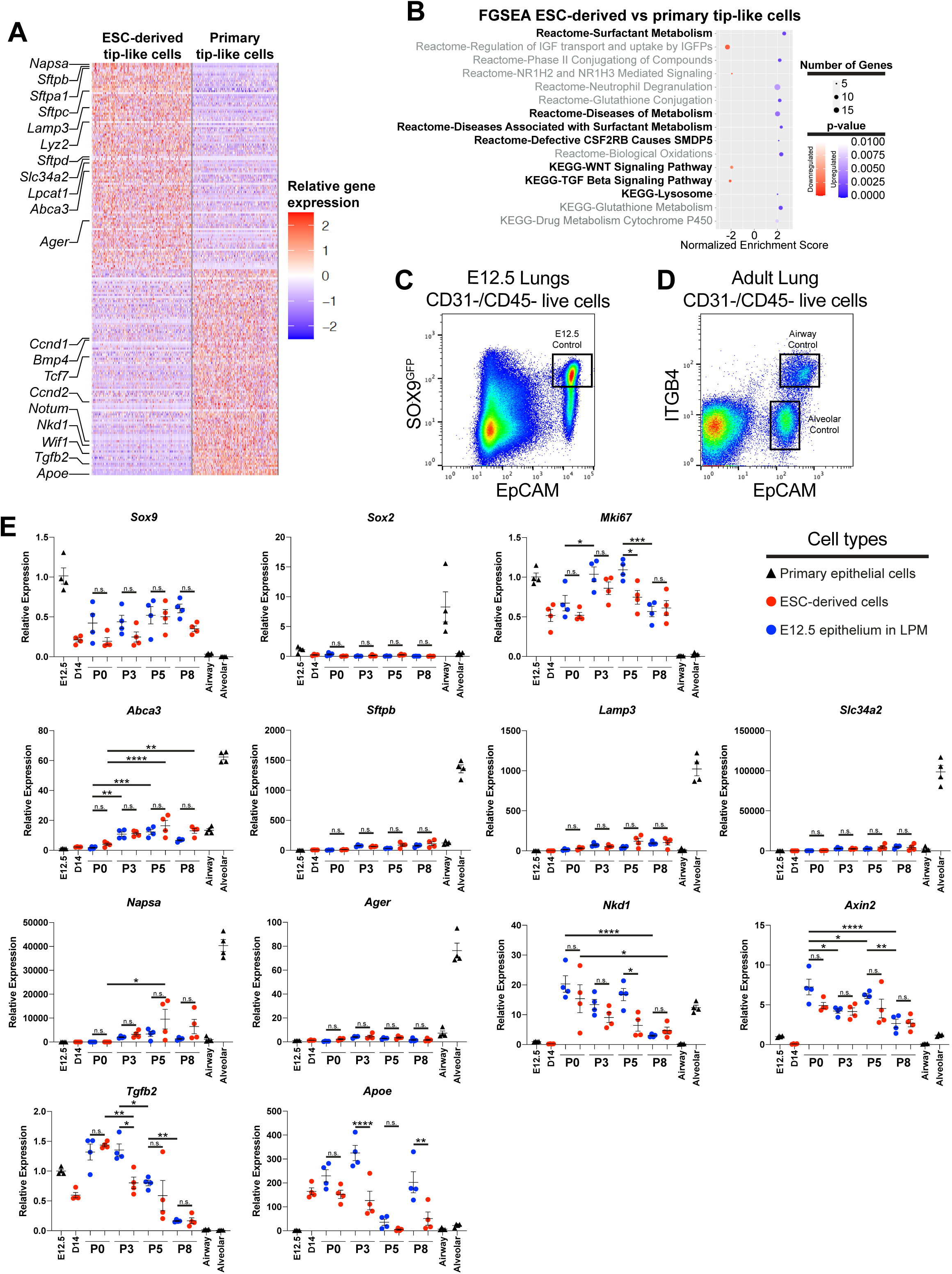
Transcriptomic Analysis of ESC-derived and Primary Tip-like Cells. (A) Row-normalized heatmap of the top 100 most up-regulated and top 100 most down- regulated genes (with adj. p-value <0.05, ordered by logFC) between ESC-derived and Primary Tip-like Cells. Annotated genes are associated with surfactant metabolism or signaling pathways identified in supplemental figure 3B. (B) FGSEA-identified Reactome and KEGG categories that are differentially regulated between ESC-derived and primary tip-like cells. The highlighted categories contain genes associated with AT2 function or prominent signaling pathways in lung development. Shown here are the 15 categories with the lowest p-value, a full list can be found in Table S2. (C) Representative gating used to collect E12.5 SOX9+ epithelial cells as a primary control for RT-qPCR. (D) Representative gating used to collect adult airway and alveolar epithelial cells as primary controls for RT-qPCR. (E) Analysis of gene expression by RT-qPCR. Primary and ESC-derived tip-like cells from multiple passages are compared against lung epithelial progenitors from day 14 of the WBRF protocol (D14) and freshly sorted lung epithelial cells from embryonic (E12.5) and adult (Airway and Alveolar) mouse lungs. n.s. not significant, * p<0.05, ** p<0.01, *** p<0.001, **** p<0.0001 by one-way ANOVA. n= 4 biological replicates. Dots with error bars = average +/- SEM.

**Supplemental Figure 4:**
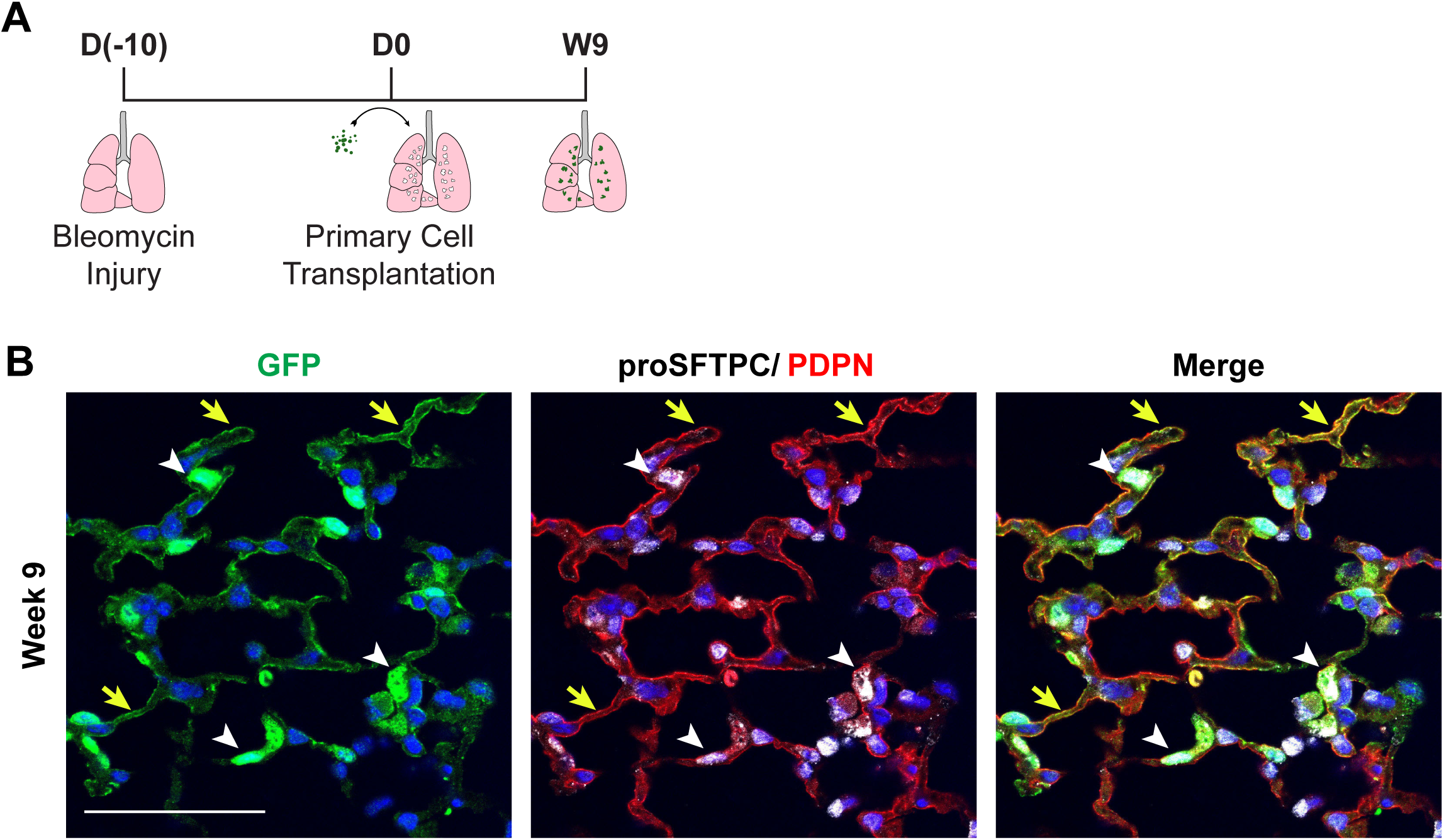
Transplantationof Primary Tip-like cells into an lmmunocompetent Recipient. (A) Schematic for transplantation of GFP+ primary cells into bleomycin injured lungs with later histological assessment of recipient lungs. (B) Representative immunofluorescence confocal microscopy of donor-derived cells at 9 weeks post-transplantation with antibodies detecting GFP, proSFTPC, and PDPN. White arrowheads indicate cuboidal proSFTPC+/GFP+ cells and yellow arrows indicate thin PDPN+/GFP+ cells. Scale bar is 50um.

**Supplemental Figure 5:**
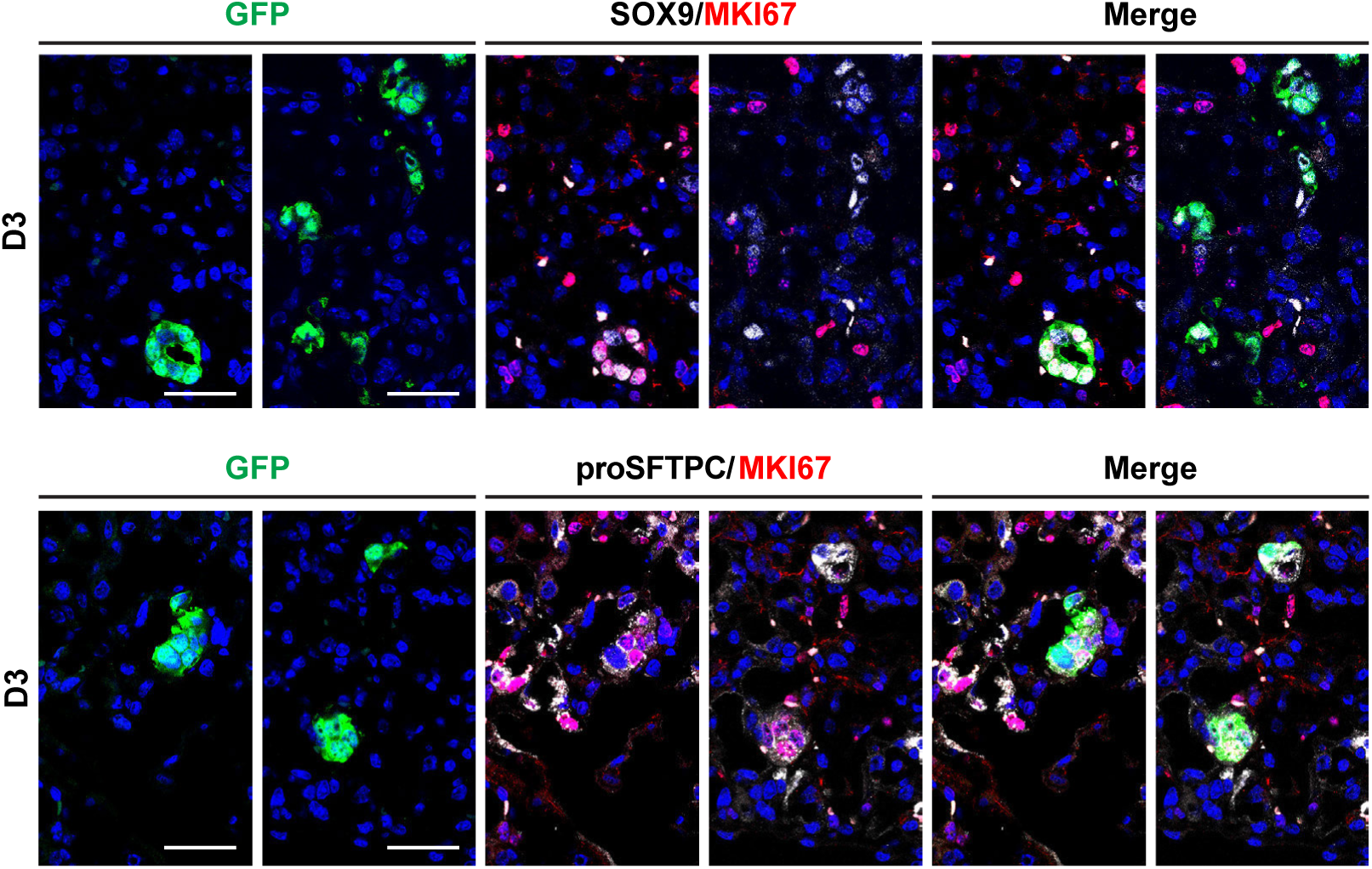
ESC-derived Tip-like Cells Maintain Proliferative Tip-like State in vivo 3 Days After Transplantation into Recipient Lungs. Representative immunofluorescence confocal microscopy of lung tissue sections containing donor-derived cells at 3 days post-transplantation. Sections stained with antibodies detecting GFP, proSFTPC, SOX9, and MKl67. Scale bars are 25um.

**Supplemental Figure 6:**
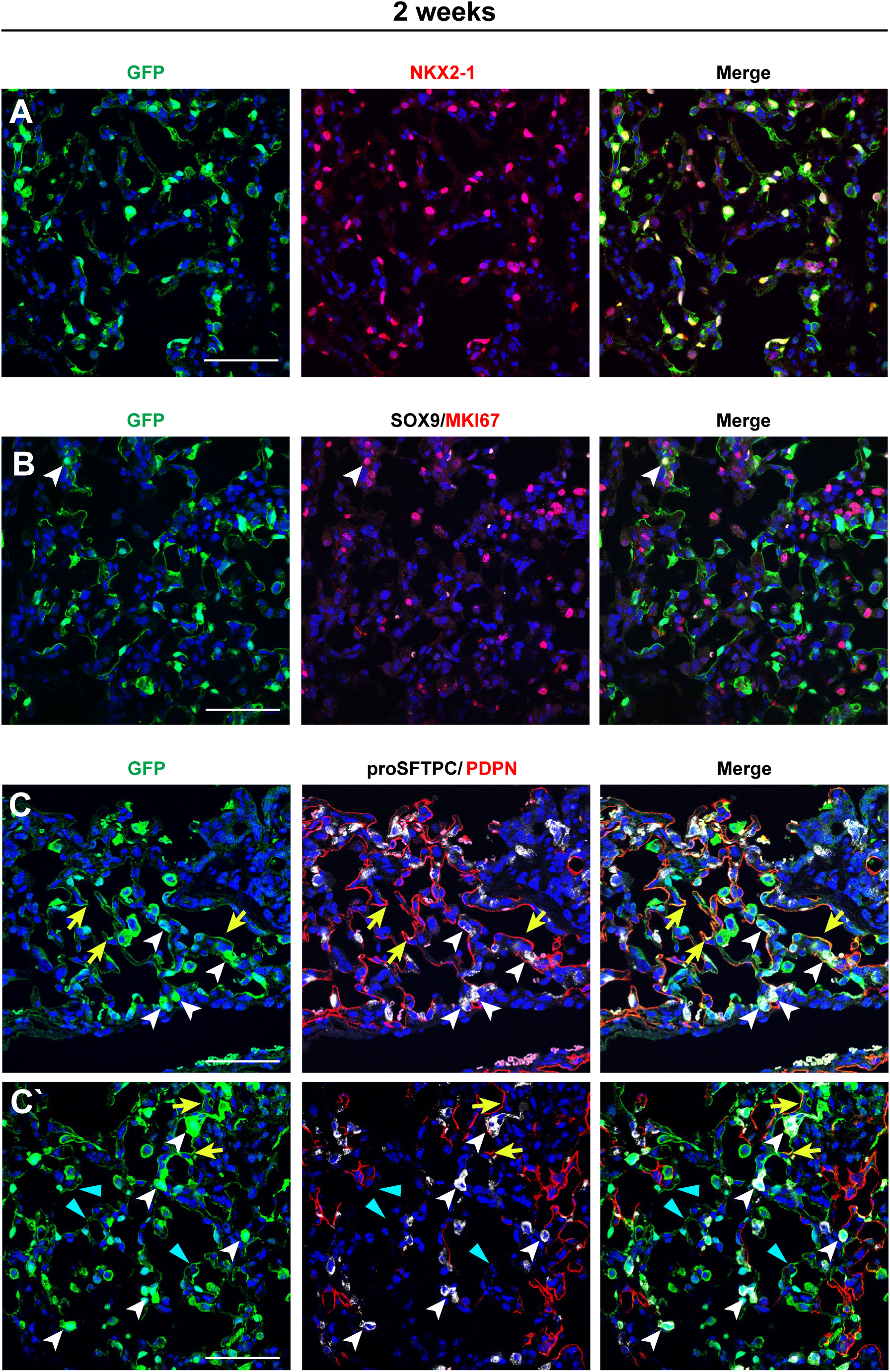
At 2 Weeks Post-transplantation into Recipient Lungs Donor-derived Cells Have Differentiated and Become Less Proliferative. (A) Representative immunofluorescence confocal microscopy of lung tissue sections indicating donor-derived cells (GFP+) at 2 weeks post-transplantation of ESC-derived tip-like cells. The majority of GFP+ donor-derived cells are NKX2-1+. Scale bars are 50um. (B) At this time point donor-derived cells are SOX9- and only a small fraction are MKI67+ (white arrowhead). Scale bars are 50um. (C) At this time point donor-derived cells include both cuboidal proSFTPC+ (white arrowheads) and thin PDPN+ (yellow arrows) donor-derived cells. Some donor-derived clusters (C’) are primarily composed of cuboidal proSFTPC+ (white arrowheads) and thin PDPN- cells (blue triangles). Scale bars are 50um.

**Supplemental Figure 7:**
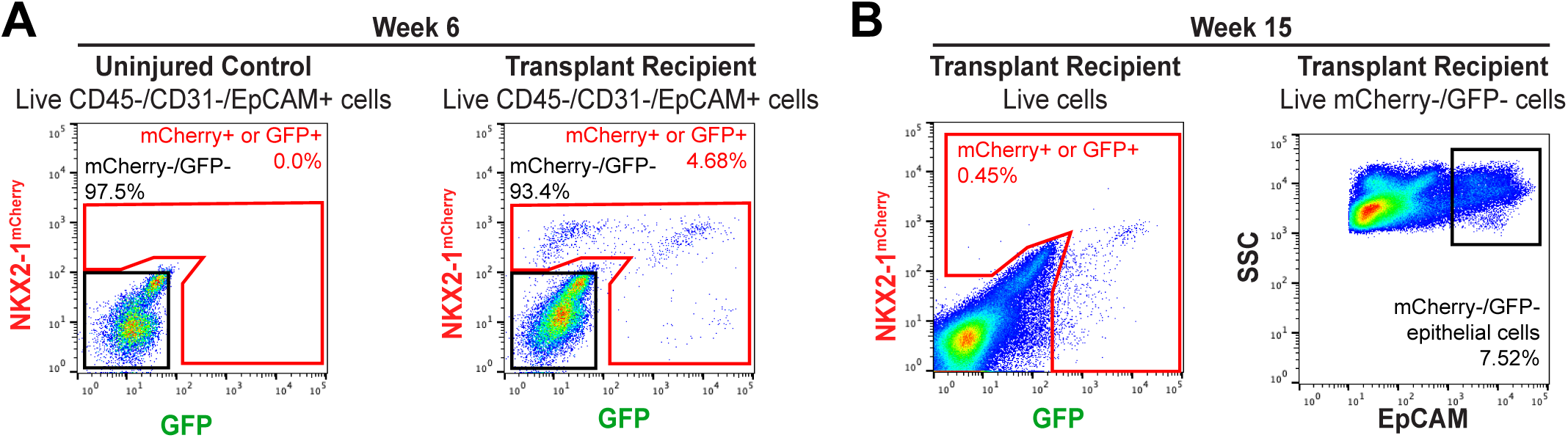
Gating for Collection of Primary Samples for scRNA-seq. (A) Gating for FAGS-based collection of endogenous (mGherry-/GFP-) and donor-derived (mGherry+ or GFP+) lung epithelial cells from an uninjured control and a transplant recipient at 6 weeks post-transplantation of ESG-derived tip-like cells. (B) Gating for FAGS-based collection of endogenous epitheilal (mGherry-/GFP-/EpGAM+) and donor-derived (mGherry+ or GFP+) cells at 15 weeks post-transplantation of ESG­ derived tip-like cells.

**Supplemental Figure 8:**
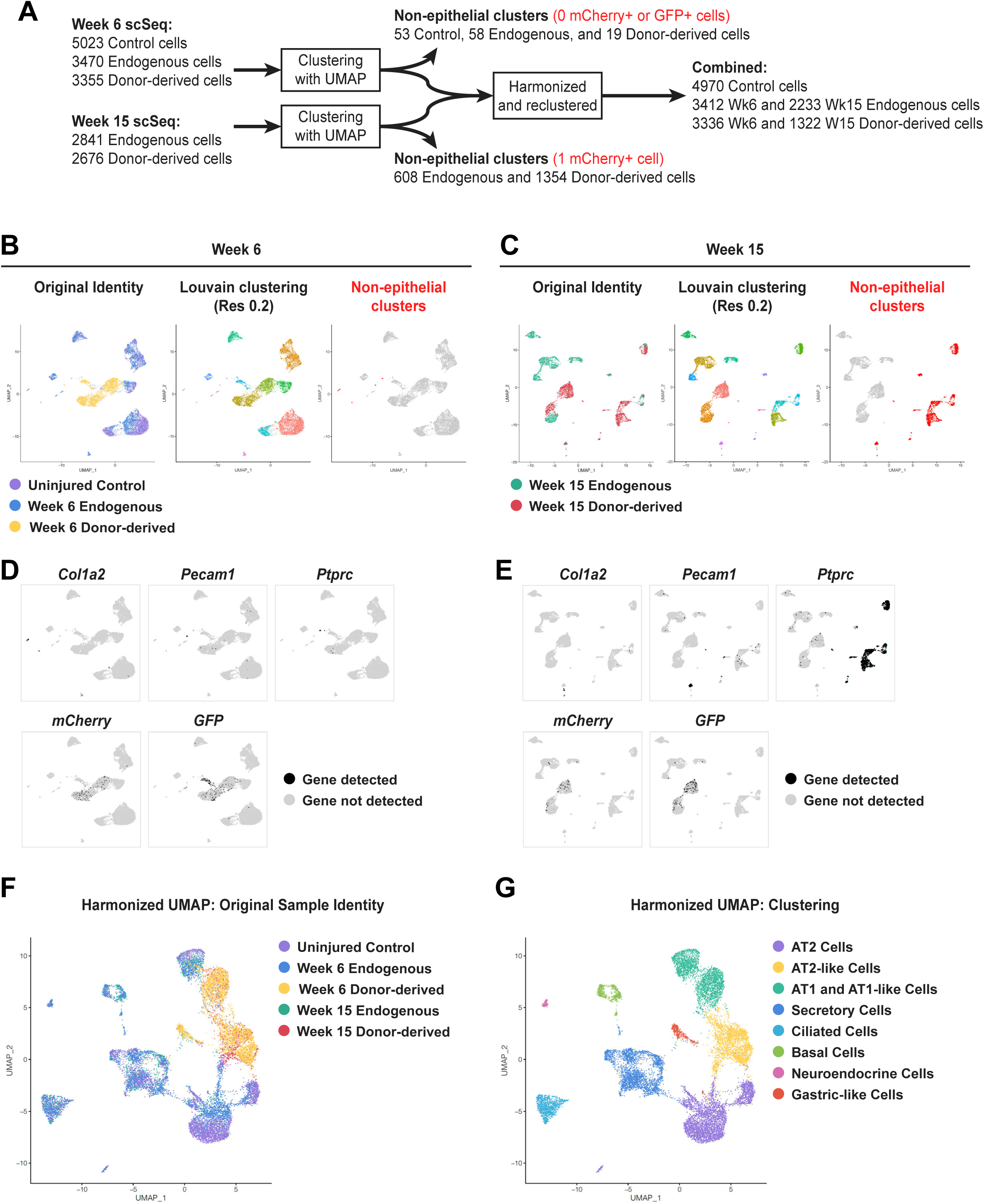
UMAP-based Analysis and Clustering of scRNA-seq Data. (A) Pipeline for analysis of scRNA-seq data including exclusion of non-epithelial lineages based on minimal presence of true donor-derived cells in these populations. (B) UMAP plots for samples collected 6 weeks post-transplantation. Plots identify sample origin, Louvain clustering, or non-epithelial cells. (C) UMAP plots for samples collected 15 weeks post-transplantation. Plots identify sample origin, Louvain clustering, or non-epithelial cells. (D) *Col1a2*, *Pecam1*, and *Ptprc* expression was used to identify non-epithelial cell clusters in samples collected 6 weeks post-transplantation. Expression of *mCherry* and *GFP* were used to verify donor-derived clusters. (E) *Col1a2*, *Pecam1*, and *Ptprc* expression was used to identify non-epithelial cell clusters in samples collected 15 weeks post-transplantation. Expression of *mCherry* and *GFP* were used to verify donor-derived clusters. (F) Epithelial cells from both timepoints were combined using harmonization to generate a single UMAP plot. (G) Cells were clustered using the Louvain algorithm followed by combining overlapping clusters. Clusters were then identified based on cell type signatures outlined in Table S3.

**Supplemental Figure 9:**
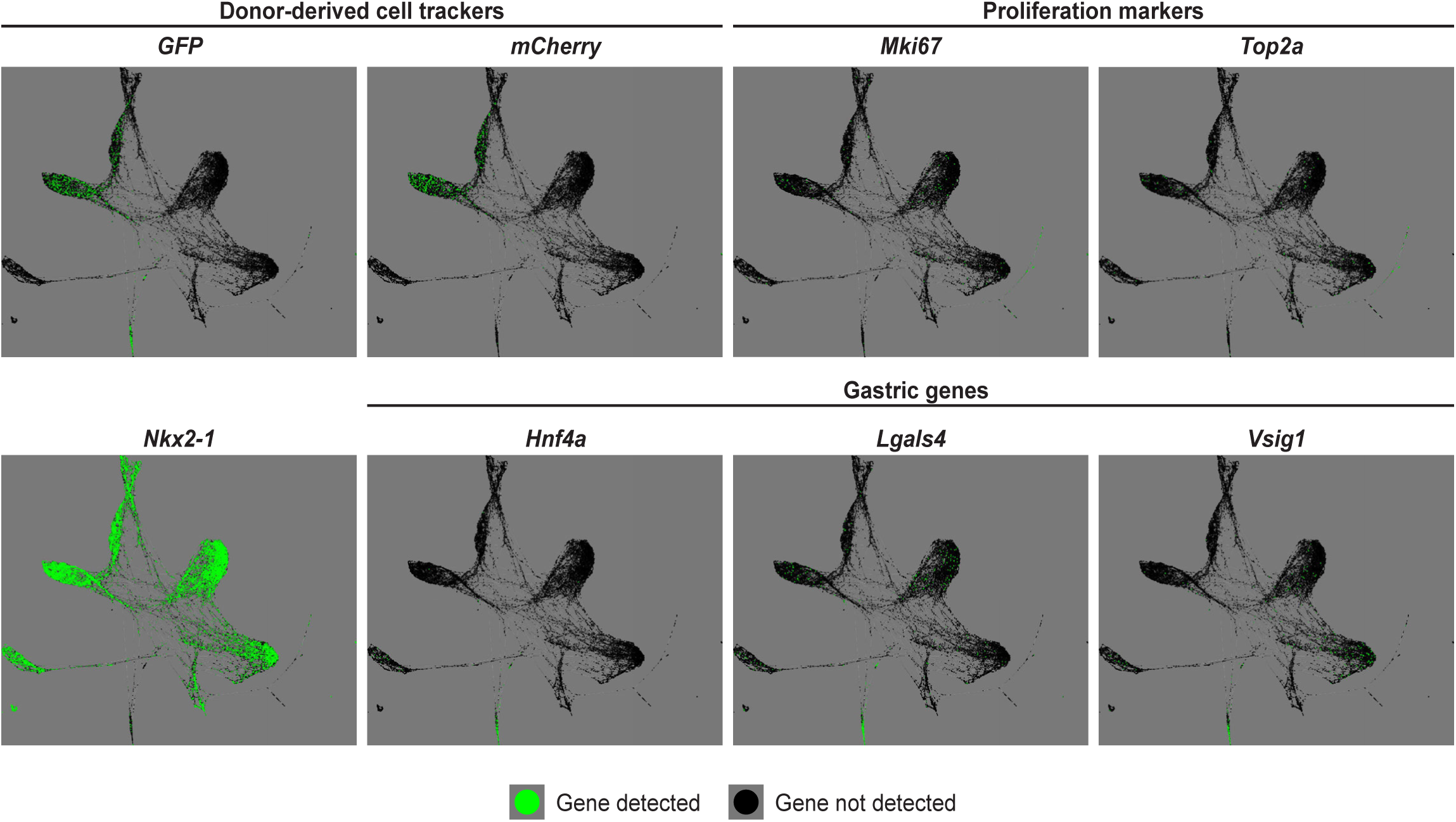
Analysis of Proliferative and Gastric Markers by scRNA-seq. SPRING plots indicating cells with detectable expression of *GFP, mCherry,* proliferation markers, *Nkx2-1,* and gastric genes in endogenous and donor-derived cells at 6 and 15 weeks post-transplantation. See figure 4 for annotation of SPRING plot by sample origin or cell type.

**Supplemental Figure 10:**
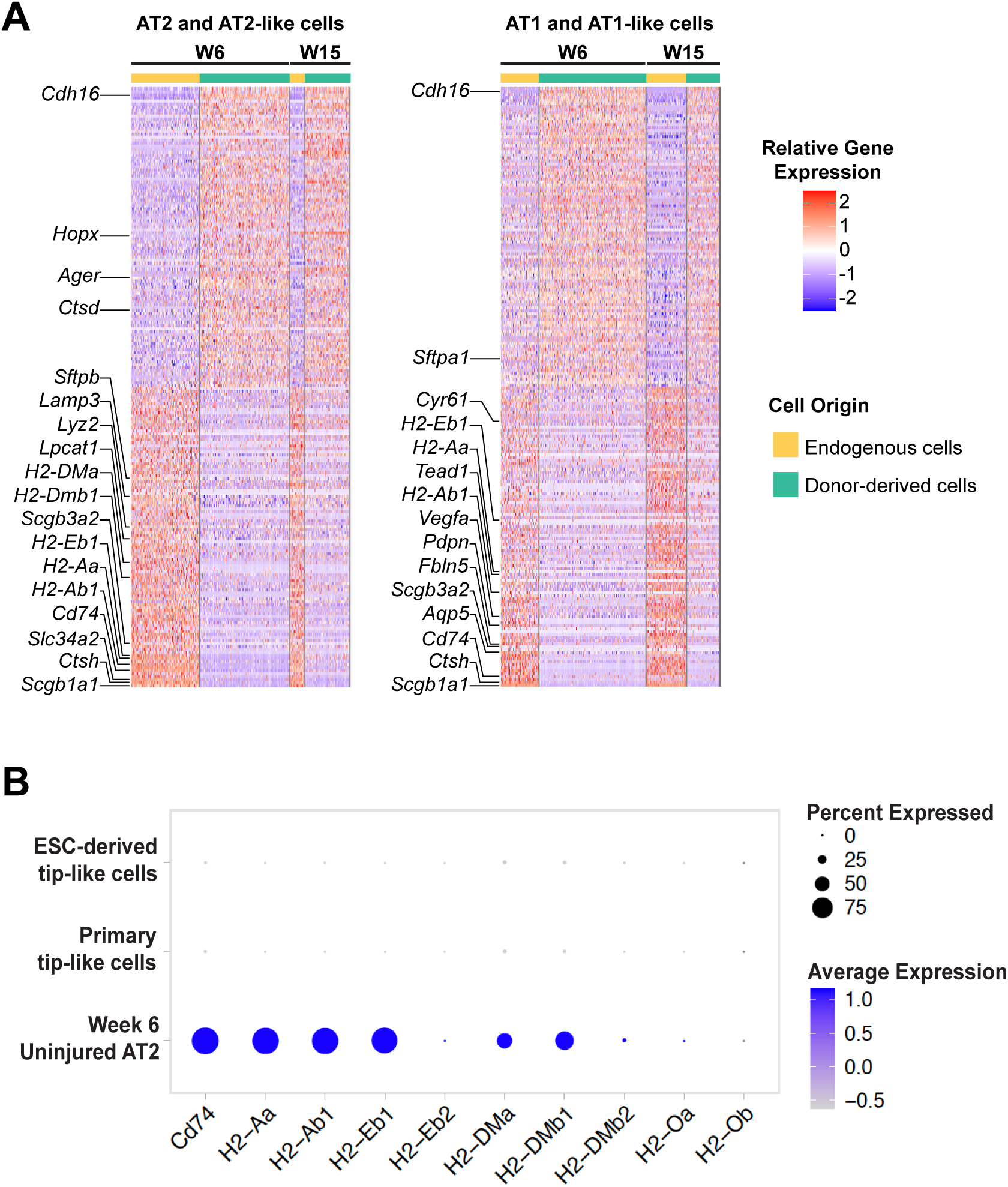
Direct Comparison of Donor-derived and Endogenous Epithelial Lineages 6-15 Weeks After Transplantation. (A) Row-normalized heatmap of the top 100 most up-regulated and top 100 most down­ regulated genes (with adj. p-value <0.05, ordered by logFC) between donor-derived and endogenous cells for both AT2-like and AT1-like lineages. Annotated genes are associated with lung epithelial lineages or MHC-11. (B) Expression of MHC-11genes in ESC-derived and primary tip-like cells compared to adult AT2 cells captured in the same experiment.

**Supplemental Figure 11:**
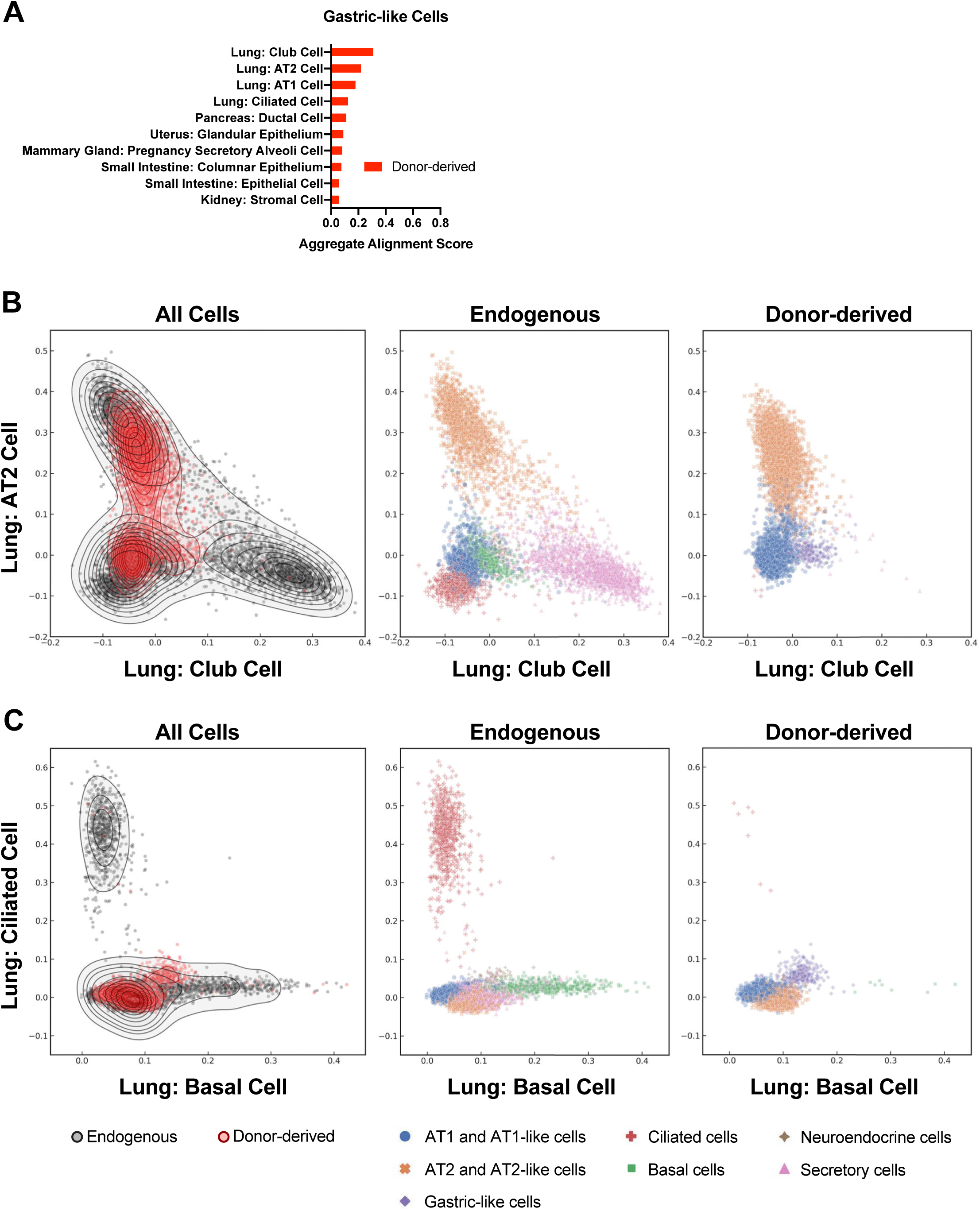
Donor-derived Cells Align Strongly With Alveolar Rather Than Airway Lineages. (A) The top ten aggregate alignment scores for donor-derived gastric-like cells. All reference cell types are from adult mice. (B) Individual alignment scores for all donor-derived and endogenous epithelial cells against reference adult lung club and AT2 cells. Cells are annotated based on sample type or cell type as determined in Figure 4. (C) Individual alignment scores for all donor-derived and endogenous epithelial cells against reference adult lung ciliated and basal cells. Cells are annotated based on sample type or cell type as determined in Figure 4.

**Supplemental Table 1: Differentially Expressed Genes Between ESC-derived and Primary Tip-like Cells**

Listed are the names and statistics (p-value, average log fold-change, proportion expressing, and adjusted p-value) for all differentially expressed genes between ESC-derived and primary tip-like cells grown in LPM as determined through single-cell RNA-seq.

**Supplemental Table 2: FGSEA for DEGs Between ESC-derived and Primary Tip-like Cells**

This table contains fast preranked gene set enrichment analysis (FGSEA) for differentially expressed genes between cultured ESC-derived and primary tip-like cells as determined by single-cell RNA-seq. Analysis was performed using REACTOME, KEGG, and C2 reference gene sets.

**Supplemental Table 3: Lung Epithelial Cell Type Gene Signatures**

This table contains lists of genes that are selectively upregulated in the predominant lung epithelial cell types and were used to generate cell type gene expression signatures used to identify cell types in scRNA-seq datasets. See the methods section for cutoffs used to generate these lists.

**Supplemental Table 4: Differentially Expressed Genes Between Donor-derived and Endogenous AT2-like Cells**

Listed are the names and statistics (p-value, average log fold-change, proportion expressing, and adjusted p-value) for all differentially expressed genes between donor-derived AT2-like cells and endogenous AT2 cells as determined through single-cell RNA-seq. Data comes from pooling two samples collected 6 or 15 weeks post-transplantation.

**Supplemental Table 5: Differentially Expressed Genes Between Donor-derived and Endogenous AT1-like Cells**

Listed are the names and statistics (p-value, average log fold-change, proportion expressing, and adjusted p-value) for all differentially expressed genes between donor-derived AT1-like cells and endogenous AT1 cells as determined through single-cell RNA-seq. Data comes from pooling two samples collected 6 or 15 weeks post-transplantation.

**Supplemental Table 6: FGSEA for DEGs Between Donor-derived and Endogenous AT2-like Cells**

This table contains FGSEA for differentially expressed genes between donor-derived AT2- like cells and endogenous AT2 cells as determined through single-cell RNA-seq. Analysis was performed using REACTOME, KEGG, and C2 reference gene sets.

**Supplemental Table 7: FGSEA for DEGs Between Donor-derived and Endogenous AT1-like Cells**

This table contains FGSEA for differentially expressed genes between donor-derived AT1- like cells and endogenous AT1 cells as determined through single-cell RNA-seq. Analysis was performed using REACTOME, KEGG, and C2 reference gene sets.

**Supplemental Table 8: Reagent Lists**

This document contains a list of all probes used for RT-qPCR, a list of all antibodies used with dilutions, and the recipes for the following medias: cSFDM, Proximal Media, Distal Media, and LPM.

## Methods

### Specification and purification of ESC-derived lung epithelial progenitors

As previously described (Ikonomou et al., 2019), Nkx2-1^mCherry^ mouse ESCs were differentiated into definitive endoderm by culturing in cSFDM for 2.5 days, trypsinizing cells to a single-cell suspension, and culturing in cSFDM supplemented with Activin A (50 ng/ml) for another 2.5 days. The resulting embryoid bodies were then grown in suspension in cSFDM supplemented with SB431542 (10 uM) and rmNoggin (100ng/ml). After one day in culture, these embryoid bodies were trypsinized and plated on six well plates coated in 100ul of Matrigel at 2e6 cells/well in cSFDM supplemented with rhBMP4 (10ng/ml), Wnt3a (100ng/ml), and Y-27632 ROCK inhibitor (10uM). Cells were fed the same media the next day and then fed daily with cSFDM supplemented with just rhBMP4 and Wnt3a. Where indicated in the text results, the media was supplemented with RA (100nM) from D6-D8 or rmFgf10 (50ng/ml) from D8-D14. On D13-14 the cells were incubated at 37C for 1 hour in 1mg/ml each of Collagenase IV and Dispase to digest the Matrigel bed. In cases where the percent of EpCAM+ cells was not being measured two slow spins (100xg) and washes were used to enrich for the undigested epithelial spheres. Epithelial spheres were then trypsinized to generate a single-cell suspension, and Nkx2-1^mCherry^+/EpCAM+ live cells were assessed by flow cytometry or sort purified for further cell culture.

### Differentiation of ESC-derived lung epithelial progenitors

In order to further differentiate D13-14 Nkx2-1^mCherry^+/EpCAM+ live cells into lung bud tip-like cells, sorted cells were resuspended in Matrigel droplets at either 200 cells/ul (LPM) or 500 cells/ul (Proximal and Distal Media) and fed every two days until collection. Media recipes can be found in table S8. In order to passage cells grown in LPM, the Matrigel droplets were incubated at 37C for 1 hour in 1mg/ml each of Collagenase IV, Dispase, and Papain with pipetting every 30 minutes. The resulting single-cell suspension was resuspended in LPM and counted on a hemocytometer.

These cells were then resuspended in Matrigel droplets as above. These cells were frozen in fetal bovine serum with 10% DMSO.

### Animal maintenance

All mouse studies were approved by the Institutional Animal Care and Use Committee of Boston University School of Medicine. All mice were purchased from Jackson Laboratory and maintained in facilities overseen by the Animal Science Center at Boston University.

### Generation of primary tip-like cells

E12.5 mouse lungs were dissected from embryos and individual lungs were digested in TrypLE Express Enzyme for 15 minutes and broken up through repeated pipetting. EpCAM+/CD45-/CD31- live cells were then sort purified and cultured at 40-200 cells/ul in LPM conditions. For P0 RT-qPCR multiple lungs were pooled to generate sufficient sample, but lines used for continued culturing were all generated from individual lungs. The sex of primary lines was determined by SRY PCR and male lines were used for all transplants.

### Reverse transcriptase quantitative PCR (RT-qPCR)

RNA was isolated according to manufacturer’s instructions using the QIAGEN miRNeasy mini kit (QIAGEN). cDNA was generated by reverse transcription of up to 100ng RNA from each sample using the Applied Biosystems High-Capacity cDNA Reverse Transcription Kit. For qPCR, technical duplicates of each of at least three biological replicates were run for 40 cycles as 10ul reactions. All primers were TaqMan probes (Table S8) from Applied Biosystems and the qPCR reactions were performed on an Applied Biosysytems Quantstudio 6 Flex. Relative gene expression was normalized to an 18S control and reported as a fold change relative to a control sample in the experiment (i.e. fold change calculated as 2^-DDCT^ based on the method of Pfaffl (Pfaffl, 2001)). Samples that were undetectable were assigned a CT value of 40 to allow for fold change calculations.

### Statistical methods

Statistical methods relevant to RT-qPCR or Flow assessment are outlined in the figure legends. Unpaired, two-tailed Student’s t tests were used for comparisons involving only two groups, while ANOVA was used when considering multiple groups.

### Immunohistochemistry

Mice were euthanized and lungs were inflation-fixed with 4% paraformaldehyde (PFA) prior to overnight fixation in 4% PFA. Cells grown in vitro were embedded in HistoGel prior to overnight fixation in 4% PFA. All samples were then dehydrated and embedded in paraffin for sectioning with a microtome (8um sections). The resulting slides were deparaffinized, blocked using normal donkey serum, and then stained with up to three primary antibodies overnight at 4C (see Key Resource Table for antibodies used). The next day slides were stained with Hoecsht and up to three secondary antibodies (see Key Resource Table for antibodies used) for 1 hour at room temperature and mounted with ProLonged Diamond Antifade Mountant. Stained slides were imaged using a Leica SP5 Confocal Microscope.

### Transplant protocol including recipient generation

Transplant recipients were syngeneic to the cells they received, unless otherwise indicated in the the text. For UBC-GFP primary tip-like cell transplants all recipients were C57BL/6J male mice. For all other transplants, recipient mice (129X1/S1) were generated by crossing 129X1/SvJ females with 129S1/SvImJ males. Recipient mice were at least 8 weeks old at the time of injury and we used both male and female 129X1/S1 mice with no clear difference in transplantation success. Ten days prior to cell transplantation mice were given 1.5U/kg bleomycin by intratracheal delivery after being anesthetized with isoflurane. On the day of transplantation donor cells were digested down to a single-cell suspension, as described above for passaging. These cells were suspended in LPM and left in a 37C incubator for 2-3 hours, with flicking every 30 minutes, to recover from digestion. At the end of this period cells were counted on a hemocytometer and resuspended in LPM (no more than 50ul/mouse with cell numbers indicated in the text results). These cell suspensions were delivered intratracheally, similar to bleomycin. Transplantation into NSG mice was performed by identical methods, except bleomycin was given 3 days prior to transplantation.

### scTOP Methods

The Python package Single-Cell Type Order Parameters (scTOP) was used to calculate alignment scores for endogenous and donor-derived cell populations. This algorithm (Yampolskaya et al., manuscript in preparation) pre-processes scRNA-seq data then finds the projection of a sample onto the space of known cell types. In brief, data was first pre-processed and normalized to reduce batch effects. To pre-process individual cells, the vector of raw RNA counts for each cell was normalized independently, 1 was added to each entry of the vector, the logarithm was taken, and the resulting data was fit onto a log-normal distribution. Next, a z-score was assigned to each gene. To do so, the vector components were first assigned a rank from least to greatest. Each rank was then divided by the total number of genes, which gave the probability that the value of a variable drawn from a normal distribution is equal to or less than that data point. Finally, the resulting percentile function was applied to a normal distribution with mean 0 and standard deviation 1. To pre-process aggregates of cells and thus find the pre- processed gene expression profile of a particular cell population, the same process is used as for individual cells, except in the very first step the average raw RNA counts of the population is used rather than the individual counts.

To find the cell type alignments for a sample, each sample’s gene expression profile was projected onto the subspace of cell types. In the following equation, the reference basis of cell types is denoted by *ξ*, which is a *p* (number of cell types in the reference basis) by *n* (number of genes) matrix. The sample is represented by a vector in gene expression space *S*. This is a vector of length *n*. Each sample in gene expression space is projected onto the hyperplane of cell type space. Thus, the sample vector in gene expression space is broken into a component that lies on the hyperplane and a component perpendicular to the hyperplane (*S*_˔_).

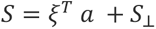

The component that lies on the hyperplane is a linear combination of all the cell type vectors. α is a *p*-length vector of the cell type components of the projected sample. The equation to find these components is:

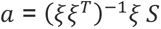

The alignment scores given in the paper represent the alignment of the sample with the indicated cell type. Individual or aggregate alignment scores were found by pre-processing and projecting the gene expression of a single cell or the average gene expression profile of a cell population, respectively.

To create the reference basis *ξ* for adult mouse cell types, we pre-processed data from the Mouse Cell Atlas (Han et al., 2018). Since the number of lung cells sampled by the Mouse Cell Atlas were relatively low for the cell types of interest, the AT1, AT2, ciliated, club, and basal cell expression profiles were taken from an uninjured control lung sample (Fig. 4), as indicated in the text. To create the reference basis from the raw count data of the atlases, the scRNA-seq counts were averaged across all cells of each cell type, then the aggregate gene expression profile was pre-processed as previously described.

### Immunogold Imaging

Mouse lung slices obtained from wild type control and mouse lung grafts containing donor GFP positive cells were removed and immediately fixed with ice cold 4% paraformaldehyde and 0.1% glutaraldehyde in 0.2M HEPES buffer, pH 7.2 for 30 min and postfixed with fresh fixative at 4 °C overnight. They were cut into 1 mm cubes, cryoprotected with PVP/sucrose (Jacob et al., 2017), frozen in liquid nitrogen and processed for freeze substitution with 0.1% glutaraldehyde, 0.1% uranyl acetate, 0.1% osmium tetroxide, and 5% DH2 in alcohol at -120 °C for 16 hours (Walther and Ziegler, 2002; Webster et al., 2008), followed by incubation with 100% alcohol at -80 °C for another 20 hours. Freeze substituted mouse lung blocks were gradually warmed up to 4 °C from -80 °C. They were washed with 100% alcohol at 4 °C, infiltrated with LR white resin, and polymerized at 60°C for 24-48 hours. Mouse lung sections were cut at 100-120 nm using a Reichert Ultrcut E ultramicrotome. Localization of GFP was visualized using a mouse monoclonal anti GFP antibody (3E6 clone; Invitrogen) and 10 nm protein A gold probes (Department of Cell Biology, University of Utrecht) described previously (Jacob et al., 2017). Electron micrographs of control and GFP positive cells were acquired using a Hitachi H-7850 TEM and an AMT BIOSPR16 TEM CCD camera at 80 kv.

### Adult lung digestion to a single-cell suspension

Mice were euthanized and perfused through injection of PBS into the right ventricle. The lungs were washed three times with 1ml of PBS administered through the trachea. Lungs were then inflated with 1.5 ml of digestion buffer (9.5U/mL Elastase, 20U/mL Collagenase, 5U/mL Dispase) followed immediately by up to 0.5ml of 1% low melt agarose and tied off with suture. These lungs were incubated in PBS on ice for 5 minutes before dissecting off lobes and placing them into 3.5 ml of digestion buffer. Lungs were incubated at 37C on a rocker for 40 minutes before being dissociated with frequent pipetting using a 10ml pipette. Cells were passed through 70um and then 40um cell strainers. If necessary, red cell lysis buffer was used to remove red blood cells. Cells were then resuspended in FACS buffer (2% FBS in PBS) and stained as described below.

### Primary cell coculture assay

Coculture protocol was used as previously described (Barkauskas et al., 2013). Pdgfra^nGFP^ animals were used to sort lung fibroblasts for cocultures (Hamilton et al., 2003). GFP+ donor-derived cells and GFP- endogenous cells were sorted from a transplant recipient at least 6 weeks after transplantation. In both cases lungs were digested as described above, but with a different digestion buffer (4U/mL Elastase, 400U/mL Collagenase, 5U/mL Dispase) and no agarose. Donor-derived or endogenous epithelia (5,000 cells) were then cultured with Pdgfra^nGFP^+ fibroblasts at a 1:20 ratio in 1:1 growth factor reduced 3D Matrigel with MTEC-plus medium (Rock et al., 2009). 90ul of matrigel and cell suspension was added to a 24-well 0.4-μm transwell insert (Falcon). Cells were cultured in MTEC-plus medium for 21 days. The resulting organoids were fixed with 4% neutral PFA and embedded in paraffin. Sections (7um thickness) were stained and imaged using standard immunofluorescence confocal microscopy protocols, as detailed above.

### Flow cytometry analysis and fluorescence-activated cell sorting (FACS)

Single-cell suspensions were prepared as described for passaging (in vitro samples) or lung digestion. When necessary, cells were stained with conjugated antibodies (see Table S8 for list of antibodies used) for 30 minutes in FACS buffer (2% FBS in PBS) at 4C and then resuspended in FACS buffer with 1:100 DRAQ7 (live/dead stain). FACS was performed on either a Beckman Coulter MoFLo Astrios or BD FACSARIA II SORP. Flow analysis was performed on a Beckman Coulter MoFLo Astrios, BD LSR II SORP, or Stratedigm S1000EXi. Resulting plots were further analyzed using FlowJo v10.7.1.

### Cell type signatures

Cell type signatures were used to identify lung epithelial cell types in scRNA-seq. Cell types were originally identified in two separate uninjured wild-type lung data sets (Control [GSM606035] from GSE200884 and Huang Protocol Epithelial Cells [GSM6046033] from GSE200883) based on Louvain clustering and known cell type markers. We then identified genes that were upregulated in a specific cell relative to all other lung epithelial cells. For cell types identified in both data sets (AT1, AT2, secretory, and ciliated cells) we took all genes that were in the 60 most enriched genes (by z score) for both data sets. For cell types found only in one data set (basal and neuroendocrine cells) we took the 20 most enriched genes (by z score) for that data set (Table S3).

### Single-cell RNA-sequencing

Single-cell suspensions were prepared and FACS purified on a Beckman Coulter MoFlo Astrios cell sorter as described above to collect live cell populations described in the results section. Single-cell RNA-sequencing was performed using the Chromium Single Cell 3′ system (10X Genomics) at the Single Cell Sequencing Core at Boston University Medical Center according to the manufacturer’s instructions (10X Genomics). The resulting samples were demultiplexed using Cell Ranger and mapped using STARsolo to the GRCm38 mouse genome reference extended with GFP and mCherry transcripts. Downstream analysis and quality controls were performed on Seurat v3.2.3 (Butler et al., 2018). We excluded from analysis cell doublets, cells containing more than 15% of mitochondrial RNA reads and cells with less than 800 genes detected (indicative of dying cells). We used SCTransform for normalization, regressing out the effect of unwanted sources of variation like that of the mitochondrial reads percentage. The Nearest Neighbors graph and Louvain clustering was based on the top 20 Principal Components. The data was plotted first using UMAP. In the case of in vivo samples, UMAP and Louvain clustering was used to identify and remove non-epithelial cells, as detailed in figure S8. Samples from distinct runs were then combined using harmonization (Korsunsky et al., 2019), and again clustered using Louvain clustering. Clusters that contained fewer than 10 cells and/or overlapped other clusters were combined with overlapping clusters to avoid the multiplicity of clusters that aren’t biologically relevant. In order to facilitate cluster annotation, cell cycle and other molecular signature enrichment were scored using the method described in Tirosh et al. (Tirosh et al., 2016). Differential expression tests were run using MAST (Finak et al., 2015), with prior gene filters to reduce the burden of multiple test corrections (min.pct = 0.25, logfc.threshold = 0.25). DEG heatmaps included a designated number of both up- and down-regulated genes with the greatest log fold change that have an adjusted p value less than 0.05. FGSEA was performed using REACTOME, KEGG, and C2 reference gene sets. When FGSEA was graphically visualized (Fig. S3B), only the 15 REACTOME or KEGG gene sets with the lowest p value were visualized to reduce visualization of redundant gene sets.

### Data and code availability

The scRNA-seq data discussed in this publication have been deposited in NCBI’s Gene Expression Omnibus (Edgar et al., 2002) and are accessible through GEO Series accession numbers GSE200886, GSE200883, GSE200884, and GSE200885 and will also be available on the Kotton Lab’s Bioinformatics Portal at http://www.kottonlab.com. The code for scTOP can be found at https://github.com/Emergent-Behaviors-in-Biology/scTOP, with the scripts used for this paper in the “Kotton” branch.

## Acknowledgements

We thank Brian R. Tilton and the entire BUSM Flow Cytometry Core for their technical assistance and guidance. We would to like to thank the entire Kotton Lab and Center for Regenerative Medicine of Boston University and Boston Medical Center for their support and suggestions throughout the course of this research. We would like to thank Carla Kim for insightful manuscript input and Jeff Whitsett for his guidance and support. This work was supported by NIH grants T32HL007035, F32HL149263, an NHLBI Progenitor Cell Translational Consortium (PCTC) Jump Start Award, and a Boston University Kilachand Multicellular Design Program Accelerator Grant to M.J.H; and U01HL134745, U01HL134766, U01HL148692, and R01HL095993 to D.N. K. C.-L.N. was supported by the PCTC consortium (U01HL134745).

## Author Contributions

M.J.H. and D.N.K. designed the project and wrote the paper. M.J.H., L.M., and M.M. performed directed differentiations and associated analysis. M.J.H. and B.R.T. performed co-culture experiments. C.-L.N. performed immunogold staining and EM imaging. M.J.H. and J.H. performed scRNA-seq experiments. J.L.-V., F.W., and C.V.-M performed all bioinformatic processing and analysis of scRNA-seq datasets. M.Y. and P.M. designed and performed scTOP analysis. D.N.K. and P.M. supervised research.

